# A Neural Signature of the Bias Towards Self-Focus

**DOI:** 10.1101/2024.02.01.578262

**Authors:** Danika Geisler, Meghan L. Meyer

**Author notes:** Corresponding Author: Meghan L. Meyer Department of Psychology Columbia University.

## Abstract

People are remarkably self-focused, disproportionately choosing to think about themselves relative to other topics. Self-focus can be adaptive, helping individuals fulfill their needs. It can also go haywire, with maladaptive self-focus a risk and maintenance factor for internalizing disorders like depression. Yet, the drive to focus on the self remains to be fully characterized. We discovered a brain state that when spontaneously brought online during a quick mental break predicts the desire to focus on oneself just a few seconds later. In Study 1, we identified a default network neural signature from pre-trial activity that predicts multiple indicators of self-focus within our sample. In Study 2, we applied our neural signature to independent resting-state data from the Human Connectome project. We found that individuals who score high on internalizing, a form of maladaptive self-focus, similarly move in-and-out of this pattern during rest, suggesting a systematic trajectory towards self-focused thought. This is the first work to “decode” the bias to focus on the self and paves the way towards stopping maladaptive self-focus in its course.

**Significance Statement:** Self-help aisles in bookstores, the popularity of self-care culture, curating identities on social media, and the mental health crisis surrounding the rise in depression and anxiety are all symptoms of modern life’s emphasis on the self. Why are people so preoccupied with themselves? Our results suggest self-focus may emerge spontaneously because of the brain state people enter in the default network as soon as they have a mental break. The brain state participants entered in the first few seconds of rest could be used to decode whether they next chose to focus on themselves, as well as subjective and neural markers of self-focus. Moreover, internalizing, a maladaptive form of self-focus, corresponded with systematic timing of the pre-self pattern during a resting state scan, perhaps shaping the timecourse of spontaneous, self-focused thought.

---

Writer David Foster Wallace once referred to the self as “our default setting” (Wallace, 2009). Psychological data support his view. Across cultures, people disproportionately think about themselves while mind wandering (Andrews-Hanna et al., 2010; Ruby et al., 2013; Song & Wang, 2012). They are also more likely to remember and communicate self-relevant information than information unrelated to the self (Bischoping, 1993; Dunbar et al., 1997; Emler, 1990, 1994; Landis & Burtt, 1924; Naaman et al., 2010; Symons & Johnson, 1997). Even when people try to take the perspective of someone dissimilar to themselves, much of the time they still end up projecting their own point of view (Epley et al., 2004; Tamir & Mitchell, 2013). While self-focus is necessary and positive in some forms—such as detecting our personal or social needs—in its most pernicious forms, self-focus is a risk and maintenance factor for internalizing disorders such as depression (Ingram, 1990; Just & Alloy, 1997; Kuehner & Weber, 1999; Nolen-Hoeksema, 1991, 2000; Nolen-Hoeksema et al., 1994). Identifying what drives the bias towards focusing on ourselves would pave the way towards stopping maladaptive self-focus, in turn preventing its negative downstream consequences.

Yet, the processes that generate the bias towards self-focus remain to be fully determined. What drives the urge to focus on ourselves? Insight may stem from the fact that the same parts of the brain associated with self-reflection also spontaneously engage by default whenever participants briefly pause and take a mental break. Extensive research on “the self” implicates the medial prefrontal cortex, Brodmann Area 10 (MPFC/BA10), in self-reflection (Denny et al., 2012). A separate line of research consistently shows that MPFC/BA10 is a key node of the default network, known to robustly engage “by default”, without the presence of any stimuli or experimental task (Raichle, 2015). Specifically, the default network shows stronger neural activity while passively resting relative to many experimental, cognitive tasks. Here, we asked whether the brain state people enter in the MPFC/BA10, and default network more generally, as soon as they have a mental break biases them towards next wanting to focus on themselves.

Although to date this question has not yet been answered, there are clues from the task and resting-state fMRI literatures consistent with the possibility. Task-based fMRI research on the self predominantly instructs participants to reflect on themselves (e.g., assess their own personality) or someone else (e.g., assess a close friend or well-known public figure’s personality) and demonstrates preferential increases in MPFC/BA10 *while* participants self-reflect (Denny et al., 2012). Another line of task-based research demonstrates elevated activity in MPFC/BA10 and self-reported self-focus during repetitive tasks that do not instruct participants to self-reflect (Andrews-Hanna et al., 2010; Mason et al., 2007; McKiernan et al., 2006; Smallwood & Schooler, 2015). Both areas of research have been highly generative but fail to identify neural processes that may set self-focus in motion. Meeting that goal would require examining neural activity *before* people spontaneously focus on themselves and assessing whether these neural patterns temporally predict wanting to focus on oneself. This alternative approach fits with predictive coding accounts of brain function, which broadly suggest endogenous, default brain states predict subsequent perception and cognition (Friston, 2018; Huang & Rao, 2011; Hutchinson & Barrett, 2019). For example, pre-stimulus fusiform gyrus activity predicts which of two competing visual stimuli is perceived (Hsieh et al., 3/2012) and pre-stimulus hippocampal patterns shape stimulus encoding and memory (Addante et al., 2015; Aly & Turk-Browne, 2016; Park & Rugg, 2010).

In the resting-state fMRI literature, many scholars have speculated that engaging MPFC/BA10, and the default network more generally, during stimulus-free rest may reflect some form of self-referential processing in the scanner (Raichle, 2015; Scalabrini et al., 2022). That said, while past work has shown greater resting state functional connectivity (i.e., timecourse correlations between brain regions) between MPFC/BA10 and other default network regions correlates with self-focus (Feurer et al., 2021; Kucyi et al., 2014), the direction of this relationship is unknown. Moreover, functional connectivity is too coarse a metric to parse the underlying mental representations considered during rest. Thus, linking functional connectivity patterns to self-focus relies on and falls prey to the problems posed by reverse inference (Poldrack, 2006; e.g., assuming that if you see its presence it must reflect the psychological construct it is implicated in). Advances in multivariate pattern similarity analysis may help solve this issue, as the approach allows researchers to characterize the fine-grained neural patterns reflecting specific mental representations (Haxby et al., 2014; Kriegeskorte et al., 2008). If it is possible to identify a multivariate pattern in default network regions reflecting the desire to focus on the self, this pattern could be applied to resting-state data to quantify whether endogenously engaging it predicts the bias towards self-focus.

## Study 1: Identifying a Neural Signature that Predicts Self-Focus

In Study 1, we took a multipronged approach to determine if there is a multivariate brain state spontaneously brought online during momentary rest that nudges self-focused thought, allowing us to identify a neural signature of the bias towards self-focus. First, we developed a paradigm designed to behaviorally measure the bias towards self-focus. In this paradigm, participants believed they were deciding which kinds of experimental trials they would get in a separate task that would immediately follow. The structure of the decision task was as follows: it started with a brief (∼4.5 secs) pre-trial rest period followed by a trial in which participants choose if they would later like to think about themselves, a close other (i.e., nominated friend), or a well-known other (i.e., President Biden) across multiple dimensions assessed separately (i.e., personality and physical traits; social roles; preferences; past and future). Each choice was followed by an attention reorienting trial (i.e., shape matching) to help ensure participants cleared their mind before the next pre-trial rest period (see Fig. 1A). Participants completed this task while undergoing functional magnetic resonance imaging (fMRI) and their behavioral responses indeed reflected a bias towards self-focus: they disproportionately chose to think about themselves (vs. the close and well-known others) and were preferentially faster to do so (see Results). Critical to the question of what makes us want to focus on ourselves, we assessed multivariate neural responses to each pre-trial rest phase of this task and tested if it predicted wanting to focus on the self (vs. others) on the immediately following trial. To this end, we trained support vector machine (SVM) classifiers in MPFC/BA10, and the default network more generally, to determine if multivariate patterns during pre-trial rest predict the immediately following choice to think about the self (vs. others). This approach allowed us to determine a neural pattern that can be used to “decode” the desire to think about the self before a behavioral response is made. We call this neural signature the “pre-self pattern.”

**Fig. 1.**
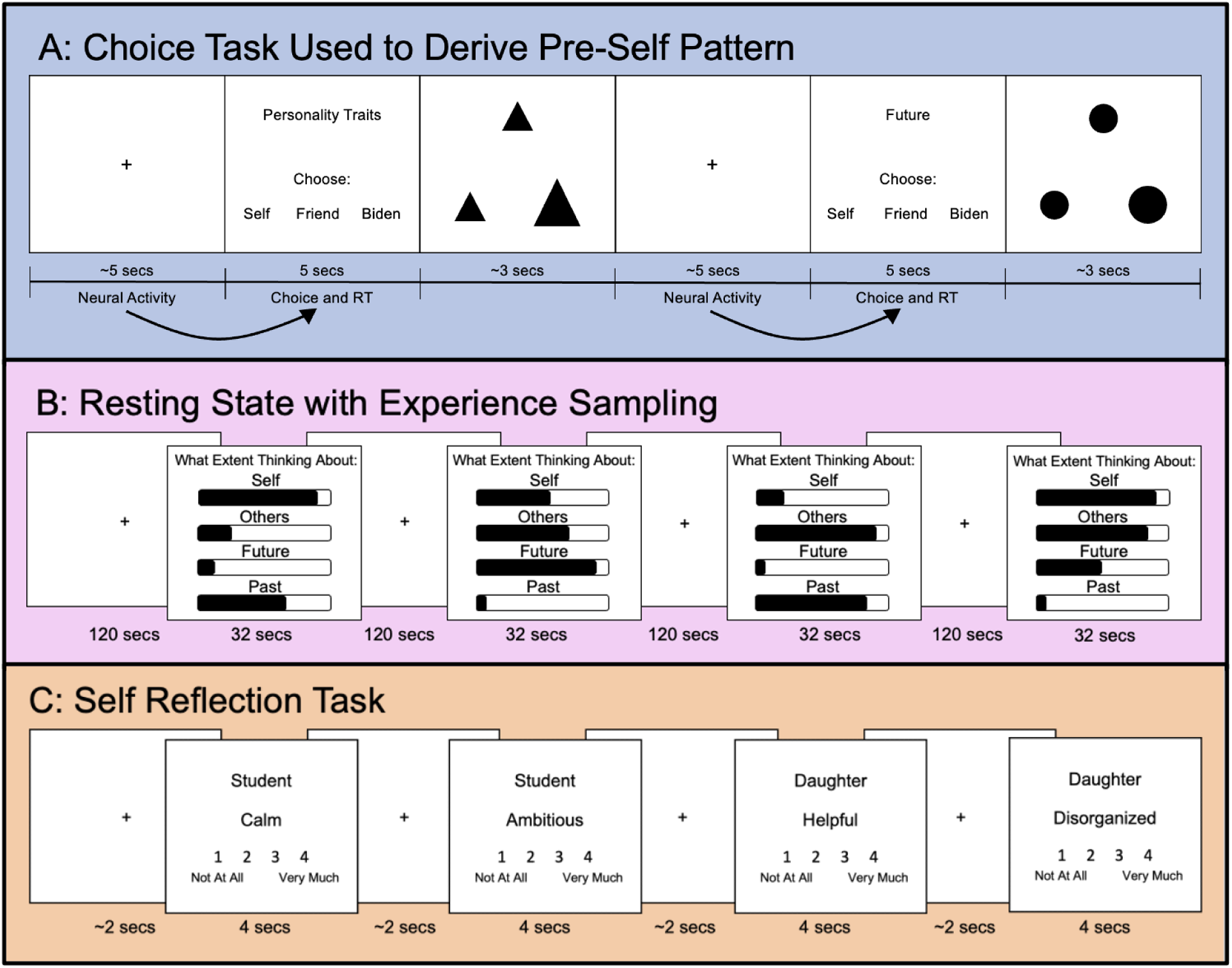
A) The choice task (used to derive the pre-self pattern) included 3 parts that repeated. First was a pre-trial jittered rest period (2.5–6 sec, mean = 4.5 sec); second was the choice activity where participants chose who (themselves, a designated friend, or Biden) they wanted to think about in a later task (5 sec); third was a shape matching task aimed at moving participants’ minds off their last choice. These three parts repeated 54 times in run 1 and 54 times in run 2 for a total of 108 trials. Our analysis examined neural activity during the pre-trial jittered rest in relation to the subsequent task choice and response time. B) The first fMRI run that the participants completed was an 8 minute long resting state scan. The 8 minutes were broken up into four, 2 minute long sections. After each 2 minute section, participants had 32 seconds to rate the extent to which they were thinking about themselves, others, the future, and the past. These ratings were made on a continuous scale with ‘not at all’ on one end, ‘completely’ on the other end, and ‘somewhat’ in the middle of the scale. C) The final task participants completed was a typical self reflection task. In this task participants were instructed to rate how well an adjective describes their personality in different social roles (Friend, Student, Significant Other, Son/Daughter, and Worker) on a scale from 1 to 4 using button boxes. We used the Big 5 list of 100 adjectives (Goldberg, 1993a). The rating trials were 4 seconds long and the jittered rest was 1–3 sec, mean = 2 sec. There were two runs of 101 trials each, with a total of 202 trials.

For a result to count as a “neural signature”, it must be generalizable – it should be able to predict self-focus in multiple ways. We therefore investigated if our pre-self pattern generalizes to predict self-focus in another context—a resting state scan. We assessed this possibility in two ways. First, in the very beginning of our experiment, participants completed an 8-minute resting state scan and rated the extent to which they were focused on themselves, others, the past, and the future every 2 minutes of the resting state scan (Fig. 1B). We took the pre-self pattern identified in the pre-trial rest period from the task described above and applied it to the resting state data. Specifically, we used multivariate pattern similarity analysis to quantify the extent to which the pre-self pattern was present during each second of the two minutes preceding participants’ self-reports during the resting state scan. The approach taken here is similar to “reinstatement” analysis from the memory consolidation literature, in which researchers assess the extent to which multivariate patterns from encoding reappear during post-encoding rest (Schapiro et al., 2018; Tambini & Davachi, 2019). Here, we refer to the approach as “instatement” analysis because we are investigating if a pre-self pattern (rather than a pattern for something previously encoded) spontaneously appears during a baseline resting state scan. We reasoned that if we discovered a neural signature that predicts self-focus, then the strength of its presence during a resting state scan should preferentially predict self-reported self-focus, too.

Second, a neural signature that temporally predicts self-focus should also temporally predict the neural pattern that captures active self-reflection. To examine this, we capitalized on a separate task completed by our participants. At the end of this experiment, participants completed a traditional self-reflection task, in which they rated their own personality traits (Fig. 1C). This approach has been used numerous times in the social neuroscience literature to capture active self-reflection (Craik et al., 1999; Fossati et al., 2003; Lieberman et al., 2004; Rogers et al., 1977; Wagner et al., 2012, 2019). For each participant, we derived their multivariate pattern (in MPFC/BA10 and default network more generally) associated with active, instructed self-reflection. We then went back to our resting state data and examined whether the strength of the pre-self pattern temporally predicted the presence of the active self-reflection pattern (and not vice versa). Such a result would offer further support that the pre-self pattern predicts self-focus. More broadly, positive results from our multi-pronged approach would provide robust evidence that we identified a neural signature that predicts multiple characterizations of self-focus (i.e., decisions, self-report, and neural).

## Study 2: Implications for Internalizing and Out-of-Sample Testing

Moving back to the larger clinical implications of this work, self-focused thought is a risk and maintenance factor for internalizing disorders, including depression and anxiety (Ingram, 1990; Just & Alloy, 1997; Kuehner & Weber, 1999; Nolen-Hoeksema, 1991, 2000; Nolen-Hoeksema et al., 1994). If we can identify a neural signature that allows us to better understand what nudges individuals into these maladaptive states, it would pave the way towards stopping maladaptive self-focus in its course, in turn preventing its negative downstream consequences. Additionally, if we have truly identified a neural signature, we should be able to apply it to individuals from separate datasets and see that it relates to participants’ self-focus. In Study 2, we tested whether our pre-self pattern predicts outcomes related to maladaptive self-focus in data from the Human Connectome Project. The Human Connectome Project dataset includes a resting state scan and internalizing score for each participant, which reflects symptoms like anxiety and depression that are related to self-focus.

We arbitrated between two ways the presence of the pre-self pattern may predict internalizing. The first possibility is a matter of “degree”: an overall greater amount of pre-self pattern instatement during a resting state scan may predict internalizing. Consistent with this possibility, individuals with (vs. without) internalizing disorders subjectively report being more strongly focused on themselves relative to other topics (Andrews-Hanna et al., 2013; Smith & Greenberg, 1981) and are more likely to reference their personal experience in laboratory tasks (Ingram & Smith, 1984) and in naturalistic settings, such as everyday conversation (Jacobson & Anderson, 1982; Mehl, 2006). It is possible that these behavioral outcomes indicating greater self-focus occur, at least in part, because individuals prone to internalizing have a stronger, default presence of their pre-self pattern. That said, the “degree” possibility is complicated by the fact that while internalizing has been linked to the default network, the literature is mixed with respect to whether internalizing corresponds with greater or weaker default network engagement (Berman et al., 2011; Javaheripour et al., 2021; Modi et al., 2015; Tozzi et al., 2021). While internalizing clearly relates to self-focus, it is not clear that this would be due to an overall stronger engagement of the pre-self pattern in the default network.

The second possibility is a matter of “fashion”, or the way in which internalizers dynamically move in and out of the pre-self pattern over time. Recent work shows that people who are similar to one another on a psychological dimension also show similar timecourses of neural engagement, not only in response to stimuli, but also during resting state scans (Finn et al., 2018; Iyer et al., 2024; Liu et al., 2019; Qiu et al., 2024; van Baar et al., 2021). The “fashion” possibility fits with research on rumination–repetitive and recurrent negative thinking about oneself (Watkins, 2008)–which is thought to play a key role in internalizing disorders (Baez et al., 2023; du Pont et al., 2018; Watkins & Roberts, 2020). This work highlights that it is the tendency to circle back to self-focus that is characteristic of internalizing disorders. Mirroring the flow of ruminative thought, there may be a temporal structure to the presence of the pre-self pattern among individuals who score high on internalizing measures.

Results were consistent with the second possibility. Human Connectome Project participants with higher internalizing scores showed a similar pre-self pattern timecourse throughout the resting state scan. In contrast, individuals low in internalizing do not show any systematic structure to the presence of their pre-self pattern; they move into and out of the pre-self pattern idiosyncratically. These results not only show the pre-self pattern can meaningfully predict maladaptive forms of self-focus in another sample. They further identify a unique way to think about how internalizing relates to self-focus. Individuals high on internalizing similarly transition in and out of the pre-self pattern, perhaps systematically impacting the timing of their spontaneous, self-focused thought.

## Study 1: Identifying a Neural Signature that Predicts Self-Focus Methods

### Participants

Thirty-two individuals (19 identifying as female, mean age = 22.4 years, SD=4.5 years; 21=Caucasian, 9=Asian, 5=Multiracial, 3=Latina/Hispanic, 1=Pacific Islander, 1=Prefer Not To Say) completed our fMRI study. Participants were eligible to participate if they were safe for MRI scanning (e.g., no metal in their body, not pregnant, not claustrophobic). All participants completed informed consent in accordance with the university institutional review board (IRB).

### Procedures

#### Resting State Scan

The first fMRI run that the participants completed was an 8 minute resting state scan. The 8 minutes were broken up into four, 2 minute sections. After each 2 minute section, participants had 32 seconds to rate the extent to which they were thinking about themselves, others, the future, and the past. For the first six participants this rating time was 25 seconds, but it was expanded to make sure participants were able to complete all four of the ratings. This was accounted for in all subsequent analyses by ensuring that all the rating TRs were regressed out in the residual image calculation stage of analysis detailed below. These ratings were made on a continuous scale with ‘not at all’ on one end, ‘completely’ on the other end, and ‘somewhat’ in the middle of the scale. Participants utilized a trackball mouse to make their selection and a value between 1 and 5, to the 14^th^ decimal point, was recorded for each of the categories (self, other, future, and past).

#### The Choice Task: Capturing the Bias Towards Self-Focus

For the main “choice” task, designed to measure the bias towards self-focus, participants were led to believe that they were choosing the trial types they would receive in a separate task that would follow. In actuality, after completing this choice task, all participants proceeded to get the same task, described below. The choice task started with a pre-trial jittered rest period (2.5–6 sec, mean = 4.5 sec), followed by the target choice activity where participants choose who (themselves, a close other (i.e., a self-nominated friend), or a well-known other (i.e., President Biden) they wanted to think about in the later task (5 sec). This design is an adaptation of previous fMRI paradigms used to assess self-reflection (Jenkins & Mitchell, 2011; Kelley et al., 2002; Meyer & Lieberman, 2018; Zhu et al., 2007). To make the task more engaging, and to confirm default MPFC/BA10 brain states predict self-focus broadly, as opposed to self-focus specifically along a certain dimension(s), we included six different categorical dimensions such that participants made choices about who to think about along the following dimensions: social roles (e.g., being a daughter), preferences, physical traits, personality traits, future, and past. Participants made a total of 108 choices (split into two runs each lasting 11 minutes and 40 seconds), evenly distributed between the dimensions with 18 choices per dimension. After each decision of which target they would think about in the later scanner task, participants completed an attention reorienting activity in which they picked which of two bottom shapes matched the top shape (2–4.5 sec, mean = 3 sec). This attention reorienting activity served as a “mental palate cleanse,” to get participants’ minds off of their last target choice before the next jittered rest. Participants were instructed not to actively reflect on the topic and target that they selected, but instead to make the decision that first came to mind, and next focus on the shape matching activity. See Fig. 1A for a schematic of the task.

#### Self-Reflection Task

The last fMRI task that participants completed for the study was an active self-reflection task. In this task participants rated how well an adjective described their personality in different social roles (Friend, Student, Significant Other, Son/Daughter, and Worker) on a scale from 1 to 4 using button boxes. We used the Big 5 list of 100 adjectives (Goldberg, 1993b). These are adjectives evenly split among the Big 5 (agreeableness, conscientiousness, intellect, emotional stability, and surgency) as well as positive and negative valence. The social role participants reported on shifted every 10 adjectives. We used a change in text color to ensure participants noted this change. The number of adjectives from each of the Big 5, as well as the positive and negative valence, were balanced across the five roles. The rating trials were 4 seconds long and the jittered rest was 1–3 sec, mean = 2 sec. There were two runs of 101 trials each, with a total of 202 trials. We used this task here to generate, for each subject, a neural pattern of their active self-reflection. Other analyses can be run with this task to answer separate theoretical questions, but are outside of the scope of this report and will not be examined here.

### Behavioral Analysis

For the choice task designed to detect the bias towards self-focus, we first conducted a chi-square test of independence to test whether the number of selections varied for the three targets (self vs. friend vs. Biden). Then, because the test was significant, we followed up with paired-sample chi-square tests of independence to confirm the interaction is driven by more decisions for the self. These chi-square tests of independence were performed using the R package *stats (R Core Team, 2022)* to compare choice of subject (self, friend, or Biden). An analysis of variance (ANOVA) tested whether RT varied across the target factor (self vs. friend vs. Biden). Next, because the results were significant, linear mixed models using the R package *lme4* were constructed to assess how choice of target (self, friend, or Biden) affected the response time (Bates et al., 2015).

### fMRI Collection

Brain imaging took place on a Siemens Prisma 3T scanner. Functional runs were acquired using a T2*-weighted echo-planar imaging sequence (2.5-mm voxels, repetition time [TR]= 1,000 ms, time to echo [TE] = 30 ms, 2.5-mm slice thickness, field-of-view [FOV] = 24 cm, matrix = 96, flip angle = 59, and simultaneous multislice = 4). A T2-weighted structural image was acquired coplanar with the functional images (0.9-mm voxels, TR = 2,300 ms, TE = 2.32 ms, 0.9-mm slice thickness, FOV = 24 cm, matrix = 256 256, and flip angle = 8) for the purpose of aligning functional data to brain structure during preprocessing. For the fMRI tasks, sequence optimization was obtained using optimizeXGUI in MatLab (Spunt, 2016).

### fMRI Preprocessing

For the fMRI dataset we collected, results included in this manuscript come from preprocessing performed using fMRIPrep 20.2.2 (Esteban et al., 2018, 2019) (RRID:SCR_016216), which is based on Nipype 1.6.1 (K. Gorgolewski et al., 2011; K. J. Gorgolewski et al., 2018) (RRID:SCR_002502). Per recommendations from the software developers, we report the exact text generated from the boilerplate in Supplementary Materials. Briefly here, preprocessing included skull stripping, motion-correction, slice-time correction, smoothing, registration and normalization.

### Residual Image Calculation

For the main “choice task” designed to measure the bias towards self-focus, in order to examine neural activity during pre-trial jittered rest that is not contaminated by neural responses during the target choice activity or shape matching activity, we first regressed out the effects of the target choice activity and shape matching activity as well as the effects of motion. This step is in line with prior research examining pre-trial neural responses (Inagaki & Meyer, 2020; Meyer & Lieberman, 2018; Spunt et al., 2015). All analyses of the pre-trial jitter period were run on the residual images saved from those models. Specifically, residual images were calculated using python’s *nltools* (Chang et al., 2023) by modeling the choice task and shape-matching task convolved with the canonical hemodynamic response function in a general linear model. This model included nuisance regressors for the six motion parameters (x, y, and z directions and roll, pitch, and yaw rotations), each motion parameter’s derivative and square of the derivative, linear drift, and run constants. We additionally regressed out TRs in nonsteady state and TRs that exhibited spikes of motion found from global signal outliers and outliers derived from frame differencing (each 3 SDs). We used a smoothing kernel of 6 on the residual images that were used for the parametric modulation analysis but no smoothing kernel on the residual images used for the MVPA analysis given that they require the fine-grained, voxel-level detail.

Consistent with prior resting state research (Meyer et al., 2019), we also calculated residual images for the resting state scan. To examine neural activity during rest that is not biased by neural activity during the self-report rating activity, we first regressed out the effects of the self-report rating activity as well as the effects of motion. All analysis of the resting state periods were run on the residual images saved from those models. Specifically, residual images were calculated using python’s *nltools* (Chang et al., 2023) by modeling the rating task convolved with the canonical hemodynamic response function in a general linear model. This model included nuisance regressors for the six motion parameters (x, y, and z directions and roll, pitch, and yaw rotations), each motion parameter’s derivative and square of the derivative, linear drift, and run constants. We additionally regressed out TRs in nonsteady state and TRs that exhibited spikes of motion found from global signal outliers and outliers derived from frame differencing (each 3 SDs). Finally, we used a smoothing kernel of 6.

## Multi-voxel Pattern Analysis (MVPA)

### Pre-Self Pattern Generation

To determine if we can “decode” the bias towards self-focus from pre-trial rest, for the choice task, we performed multi-voxel pattern analysis on a whole brain activation map masked (separately) by MPFC/BA10, the three subsystems of the default network (the core subsystem, dMPFC subsystem, and MTL subsystem) as identified by Yeo et al. (2011), as well as an unmasked whole brain activation map. These masks were chosen because we were interested in assessing MPFC/BA10 role’s in particular, and the default network’s role more broadly in prompting us toward self-focus. The unmasked whole brain was also analyzed because its classification accuracy acts as a useful comparison. If a mask (e.g., core subsystem) has a higher classification accuracy than the whole brain, this suggests the whole brain’s classification power is disrupted by the noise created by areas outside of the mask. We again assessed classification of self versus other, where friend and Biden were combined into the other category. We focused on this self versus other classification so that there was a near equal number of trials in each group, which helps ensure an unbiased classification result.

To generate the input for the classifier we used *nltools* (Chang et al., 2023) to create a first-level model (performed on participants’ residual images) that included three regressors reflecting the jittered rest period before each target choice: a rest before self-choice regressor, a rest before friend-choice regressor, and a rest before Biden-choice regressor. A baseline Beta of self and other (combining friend and Biden) was then extracted and utilized as the input to our classifier. In other words, for each subject, there was a different Beta value map for rest periods (1) preceding self-focused decisions and (2) preceding other-focused decisions. We trained a linear SVM to discriminate a subsequent choice of self (coded as 1 in the classification) versus other (coded as -1 in the classification) with a k-fold cross-validation procedure (Hastie et al., 2008; Wager et al., 2013; Woo et al., 2014). In the statistical learning literature (Hastie et al., 2008; Yu et al., 2020), there are many types of classification algorithms, but they generally perform very similarly on problems such as the one we pursued here. SVM algorithms such as the one we used in this study are the most widely used algorithm for two-choice classification and are robust and reasonably stable in the presence of noisy features.

We computed prediction performance using a 6-fold balanced cross-validation procedure (Cohen et al., 2010; Izuma et al., 2018). We subdivided the data into 6 separate folds (5-6 participants in each group) and used all of the data except for one fold to train the model and then tested the model using the left-out fold. We then iterated over this process for every possible fold. We chose the k-folds balanced cross-validation approach, instead of a leave-one-out subject approach, because it is less susceptible to outlier participants (Vu et al., 2022).

The 6-fold balanced cross-validation procedure was completed for each of our five neural maps (MPFC/BA10, whole brain, default network core subsystem, dmPFC subsystem, and MTL subsystem) and an average classification accuracy was calculated for each. A follow up analysis was then done following the same steps with all of the ROI’s included in Yeo’s (2011) default network core subsystem (right middle temporal gyrus, right superior frontal gyrus, left middle frontal gyrus with left superior frontal gyrus, left angular gyrus, right angular gyrus with right middle temporal gyrus, and PCC) except for MPFC/BA10 as that ROI was already analyzed. Another follow up analysis was also done utilizing the same methods with four ROIs, 1) default network core subsystem minus MPFC, 2) default network core subsystem minus the PCC 3) default network core subsystem minus MPFC/BA10 and PCC, and 4) MPFC/BA10 and PCC combined. We ran statistical tests on a total of 15 classifier models over the course of this analysis. To account for issues with multiple comparisons statistical results are only reported as significant if they have *p* < .05/15 or .003.

To test the statistical significance of these results, we generated null distributions for each ROI using 10,000 permutations of a 6-fold support vector machine classification analysis. First, for each subject we relabeled their self and other neural images randomly as self or other. Then we subdivided the data into 6 separate folds (5-6 participants in each group) and used all of the data except for one fold to train the model and then tested the model using the left-out fold. We then iterated over this process for every fold. This was repeated 10,000 times and the average classification accuracy of each permutation was used to create a null distribution. Our average classification accuracy result was then compared to this null distribution using nltools (Chang et al., 2023). This process was completed for all 15 of the neural maps mentioned above (whole brain, default network core subsystem, dmPFC subsystem, MTL subsystem, right middle temporal gyrus, right superior frontal gyrus, left middle frontal gyrus with left superior frontal gyrus, left angular gyrus, right angular gyrus with right middle temporal gyrus, PCC, MPFC/ACC, default network core subsystem minus MPFC, default network core subsystem minus the PC, default network core subsystem minus MPFC/BA10 and PCC, and MPFC/BA10 and PCC combined). Because results from this analysis implicated the default network core subsystem, all subsequent analyses described below move forward with the default network core subsystem as the primary mask and follow-up analyses testing for specificity to this subsystem are run with the other Yeo default network subsystems/ROIs (2011).

### Testing Whether Pre-Self Pattern Instatement during the Resting State Scan Predicts Subjective Reports of Self-Focus

The analysis described above answers whether multivariate patterns in the default network can decode self-focused choices. Next, once we characterized the pre-self pattern from the choice task, we could apply it to our resting state data to answer our next question. Specifically, we asked if the stronger presence of the pre-self pattern during rest predicted participants’ subjective ratings of self-focus during their resting state scan. We performed an instatement analysis–a TR-to-TR pattern matching analysis with the classifier pattern generated by the MVPA analysis of the default network core subsystem. Each TR of the eight minutes of rest was masked by the Yeo default network core subsystem ROI and then correlated with the default network core subsystem pre-self pattern. These correlation values were then averaged for the two minutes of rest before each rating and z-scored. Linear mixed models using the R package *lme4* were constructed to assess how self-report of thought content (self, other, future, and past) as well as the section of rest affected the mean correlation (Bates et al., 2015). To ensure that the regressors were not introducing collinearity to the model we ran tolerance and variance inflation factors. All VIF results were well below the threshold of 5 (Kutner et al., 2005) (Self = 1.2, Other = 1.1, Future = 1.3, Past = 1.1, and Section = 1.1) and all tolerance values were over 75% (Self = 82, Other = 90, Future = 77, Past = 88, and Section = 93). To follow up on this, analyses were also run with the MVPA pre-self classification patterns generated with the dmPFC subsystem, MTL subsystem, MFPC, PCC, and whole brain.

### Self-Reflection Pattern Generation

Next, we turned to our third question: does the presence of the pre-self pattern during the resting state scan even predict the presence of an active self-reflection neural pattern a few seconds later? This would again be consistent with the idea that the pre-self pattern temporally predicts future self-focus, here “neural self-focus.” To test whether our pre-self pattern temporally predicts the presence of active self-reflection during a resting state scan, we needed to create a neural pattern reflecting active self-reflection and subsequently examine the temporal relationship between pre-self pattern instatement and instatement of self-reflection neural activity during the resting state scan. To this end, we generated neural patterns for each subject using the self-reflection task. Specifically, images were calculated by modeling the self-reflection task convolved with the canonical hemodynamic response function in a general linear model. This model included nuisance regressors for the six motion parameters (x, y, and z directions and roll, pitch, and yaw rotations), each motion parameter’s derivative and square of the derivative, linear drift, and run constants. We additionally regressed out TRs in nonsteady state and TRs that exhibited spikes of motion found from global signal outliers and outliers derived from frame differencing (each 3 SDs). Finally, we used a smoothing kernel of 6. We created a contrast of self-reflection compared to baseline and saved the resulting image for each subject to use in the subsequent instatement analysis.

### Testing the Temporal Relationships between Pre-Self and Self-Reflection Pattern Instatement During a Resting State Scan

Given that we now had the pre-self classification pattern and, for each subject, an active self-reflection neural pattern (see section above “Self-Reflection Pattern Generation”), we could test whether the presence of the pre-self pattern temporally predicted the presence of an active self-reflection neural pattern. We performed an instatement analysis–a TR-to-TR pattern matching analysis–with the classification pattern generated by the MVPA analysis of the default network core subsystem and with the active self-reflection neural pattern generated with the self-reflection task. First, we masked the active self-reflection pattern of each subject by the Yeo default network core subsystem ROI (2011). Then each TR of the eight minutes of rest was masked by the Yeo default network core subsystem ROI and correlated with the default network core subsystem pre-self pattern as well as the masked version of the active self-reflection pattern. Linear mixed models using the R package *lme4* (Bates et al., 2015) were constructed to assess if pre-self pattern correlation strength predicted active self-reflection pattern correlation strength 0-20 TRs later (See Fig. 5A for a visualization of the approach) . We chose to use this data driven approach with multiple TR lag-times to avoid arbitrarily selecting a time delay as well as to better understand the dynamic temporal relationship between the presence of the “pre-self” pattern and active self-reflection pattern.

Analyses are performed on fisher z-transformed correlation values. We tested the correlation of the two neural patterns for each subject and did find small (*r* = .08 to -.03). We therefore ran the linear mixed models with 0-20 TR delays: 1) with the strength (i.e., correlation) of the pre-self pattern predicting the strength of the active self-reflection-pattern X TRs later and 2) with the strength (i.e., correlation) of the active self-reflection pattern predicting the strength of the pre-self pattern X TRs later. This enabled us to make sure that the correlation of the two patterns was not the cause of significant temporal results. To account for the number of statistical tests performed (41), we used a bonferroni-corrected significance value of *p* < 0.00122 (i.e., .05×41). Significant results are reported here, see Supplemental Table 1 for a full list of the results for each TR lag.

## Results

### When given the choice, people prefer to think about themselves over others

In our primary fMRI task, participants believed they were choosing trials that would appear in their subsequent fMRI task. Specifically, they choose if they would later like to think about themselves, a close other (i.e., nominated friend), or a well-known other (i.e., President Biden) across multiple dimensions assessed separately (i.e., personality and physical traits; social roles; preferences; past and future), with a total of 108 choices made (See Fig. 1A). Participants’ decisions indicated a strong predisposition to default towards self-focus (Fig. 2). A chi-square test of independence demonstrated a main effect of choice on the number of trials selected (*X*^2^ (2, *N* = 3447) = 500.71, *p* < .001). Follow-up, paired sample chi-square tests of independence showed participants choose to think about themselves significantly more than a designated friend (*X*^2^ (1, *N* = 2823) = 114.69, *p* < .001) and Biden (*X*^2^ (1, *N* = 2320) = 495.34, *p* < .001). Using an analysis of variance (ANOVA) we also found a main effect of choice on response time (*F*(2, 3431.3) = 11.97, *p* < .001). Follow up linear mixed models showed that participants were faster to make decisions to think about themselves in comparison to a friend (β = 0.11, standardized β = 0.02, *t*(3429)= 4.68, *p* = .003) and Biden (β = 0.08, standardized β = 0.03, *t*(3435) = 2.93, *p* < 0.001). Notably, participants were also more likely to choose to think about their friend than Biden (*X*^2^ (1, *N* = 1751) = 144.49, *p* < .001), and were faster to choose their friend over Biden (β = 0.08, standardized β = 0.03, *t*(3435) = 2.93, and *p* < 0.001). Supplementary Materials presents detailed results from three follow-up analyses that further confirm the robustness of the bias towards self-focus. Briefly here, the bias to choose the self over others was not due to certain dimensions (e.g., choosing to think about one’s personality traits vs. future), appeared consistently across the experimental task, and was not affected by the length of the rest preceding the decision.

**Fig. 2.**
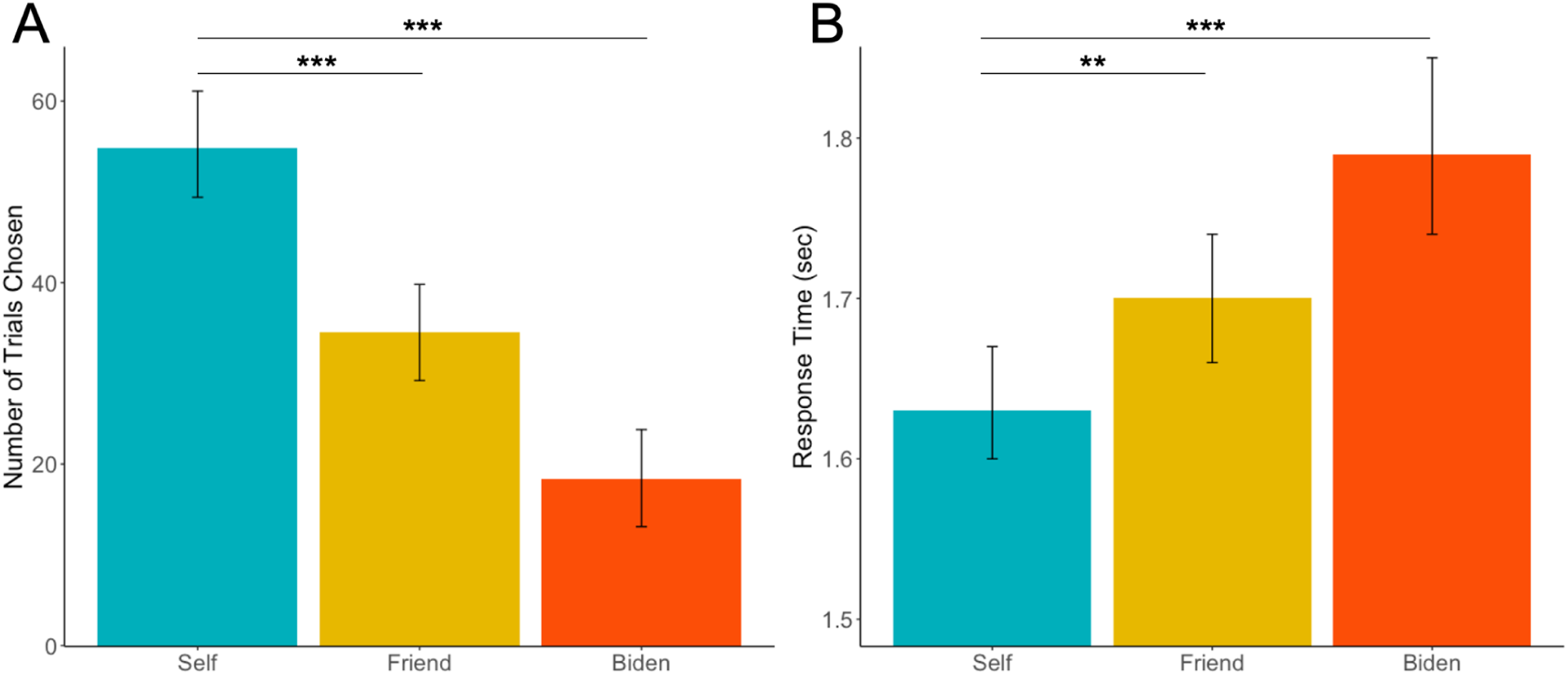
(A) Participants choose to think about themselves significantly more than their self-nominated friend (X^2^ (1, N = 2823) = 114.69, p < .001) or Biden (X^2^ (1, N = 2320) = 495.34, p < .001). (B) Participants are faster in their decisions to think about themselves in comparison to a friend (β = 0.11, standardized β = 0.02, t(3429) = 4.68, p = .003) and Biden (β = 0.08, standardized β = 0.03, t(3435) = 2.93, p < 0.001). ***indicates p < .001; **indicates p < .005.

### A neural signature that predicts self-focus: Evidence from the brief, pre-trial rest in the choice task

#### Univariate MPFC/BA10 Activity during Pre-Trial Rest Parametrically Modulates How Quickly Participants Choose to Think about the Self (vs. Others)

The primary goal of the choice task was to derive a neural signature that precedes and decodes self-focus. Meeting that goal requires using a multivariate approach to classify participants’ decisions. However, before meeting that primary goal, we first wanted to assess if we conceptually replicate the single, previous study investigating the role of pre-stimulus responses on self-referential processing (Meyer & Lieberman, 2018). To date, only one study has probed pre-stimulus neural activity in a self-reflection task, finding that, on a trial-by-trial basis, faster responses to questions assessing beliefs about one’s traits (e.g., “Am I funny?” yes/no) are preceded by stronger MPFC/BA10 activity (Meyer & Lieberman, 2018). This finding complements the present paper’s hypothesis: it shows that default MPFC/BA10 activity may help people quickly access self-knowledge. However, speed is not the same thing as desire. The prior study does not speak to whether pre-stimulus, default brain states predict wanting to focus on the self (i.e., the bias towards self-focus). We first asked whether we conceptually replicate this prior work, testing if the average amount of pre-trial activity (i.e. univariate activity) relates to task performance on a given trial—specifically, faster reaction time to self vs. other choices.

Consistent with the prior work, we performed a parametric modulation analysis in which univariate neural activity during pre-trial rest was modulated—on a trial-by-trial basis—by the speed and target choice of the next trial. Note that these and all subsequent analyses are run on the residual images from the task activation models to ensure pre-trial rest activity is not contaminated by task-evoked effects. Given that the previous work found the magnitude of MPFC/BA10 activity preferentially facilitates the speed with which participants answer questions about themselves, we probed the parametric modulation analysis here in an MPFC/BA10 region-of-interest (ROI) predefined by Yeo et al. (2011). Consistent with the prior work, faster decisions to choose the self (vs. friend and Biden) corresponded with greater mean activity in MPFC/BA10 during the previous pre-trial rest period (t(31) = -2.20, p = 0.036, Cohen’s d = -0.39). No additional clusters emerged in follow-up whole-brain parametric modulation analyses searching for any clusters whose greater activation during pre-trial rest predicted faster decisions to think about: 1) self vs other (friend and Biden), 2) friend vs. self, 3) Biden vs. self or 4) friend vs. Biden). Additionally, a follow-up, whole brain parcellation analysis with 85 ROIs defined by Yeo et al. (2011) again indicated no other regions showed a significant parametric relationship with these four contrasts (bonferroni corrected p’s > .60). Conceptually replicating prior work, the results show that faster choices to think about the self (vs. others) are preceded by greater mean MPFC/BA10 activity. For more details about these analyses, see the Parametric Modulation Methods in the Supplementary Materials.

#### Multivariate Neural Patterns in the Core Default Network Subsystem during Pre-Trial Rest Predict Decisions to Think about the Self

Next, we returned to our primary goal – to test whether we could use multivariate neural patterns during pre-trial rest to “decode” if participants subsequently wanted to think about themselves. In the decoding analysis presented here, we used the decision of target in the choice task as the outcome variable to determine if neural patterns during pre-trial rest predict the subsequent choice to think about the self. Specifically, using multi-voxel pattern analysis (MVPA) on the pre-trial rest, we trained a linear support vector machine (SVM) classifier to differentiate between subsequent self-choices vs. other-choices (friend and Biden). Friend and Biden choices were combined so there was a roughly equal number of self vs. other trial types, which helps ensure the analysis is unbiased.

To generate the input for the classifier we created a first-level model. This was performed on participants’ residual images from the task activation model, to ensure pre-trial rest activity is not contaminated by task-evoked effects. The first-level model comprised three regressors for the rest period before each trial. These regressors modeled pre-trial rest before trials in which the participant made the choice to be (1) self-focused or to be (2) other-focused. A baseline Beta value of each of the decision types–self and other–was then extracted and utilized as the input to our classifier. In other words, for each subject, there was a different Beta value map for rest periods (1) preceding self-focused decisions and (2) preceding other-focused decisions.

We first performed the analysis and validation specifically within the Yeo et al. (2011) MPFC/BA10 ROI. We computed prediction performance using the 6-fold balanced cross-validation procedure (Cohen et al., 2010; Izuma et al., 2018). Specifically, we subdivided the data into 6 separate folds (5-6 participants in each group) and used all of the data except for one-fold to train the model and then tested the model using the left-out fold. We then iterated over this process for each fold and an average classification accuracy was calculated. A null distribution was computed using 10,000 permutations of 6-fold randomized SVMs and p-values were calculated to indicate statistical significance of the predictive accuracy (see Methods). We chose the k-folds balanced cross-validation approach, instead of a leave-one-out subject approach, because it is less susceptible to outlier participants (Vu et al., 2022).

Analyses showed that distributed, pre-stimulus patterns in MPFC/BA10 predict the decision to choose to think about the self with 70% accuracy, *p* = .01. In other words, the brain state a person enters in MPFC/BA10 as soon as their mind is free from external demands predicts whether they next want to think about themselves. But is it the only part of the brain that can do this? We next assessed if MPFC/BA10 predicted self-choices better than patterns from the entire brain. The whole brain pattern during pre-trial rest was able to predict the subsequent decision to choose the self with 77% accuracy, *p* = .001. The higher accuracy for whole brain classification suggests additional brain areas contribute to the bias towards self-focus.

To determine which additional brain regions contribute to the bias towards self-focus, it is important to consider that MPFC/BA10 is one of multiple brain regions comprising the brain’s default network and that mental phenomena can arise from distributed patterns of neural activity across interacting brain regions (Bressler & Menon, 2010; Chang et al., 2015; Kragel et al., 2018). Graph analytic methods indicate that the default network is comprised of three subsystems: the core subsystem (associated with self-reflection, prospection and autobiographical memory), dorsomedial subsystem (associated with social semantic knowledge and inferring mental states), and the medial temporal lobe subsystem (associated with episodic memory, simulation, and relational processing (Andrews-Hanna et al., 2010; Buckner & DiNicola, 10/2019; Yeo et al., 2011). As alluded to in its name, the core subsystem is the primary default network subsystem, and MPFC/BA10 is a key node. Thus, while MPFC/BA10 is the region most reliably associated with self-reflection (Lieberman et al., 04/2019), it is possible that distributed patterns of neural activity in the core default network subsystem, including MPFC/BA10, may contribute to the bias towards self-focus.

We therefore repeated the steps for (SVM) classification on a whole-brain activation map masked by the three subsystems of the default network (core subsystem, dMPFC subsystem, and MTL subsystem) as mapped by Yeo et al. (2011). We found multivariate patterns in the core subsystem during pre-trial rest periods predicted the decision to choose to think about the self with 83% accuracy, *p* < .001 (Fig. 3). The dMPFC subsystem (48% accuracy, *p* = .61) and MTL subsystem (59% accuracy, *p* = .17) both were not able to classify results above chance. These results demonstrate that distributed patterns in the default network core subsystem during pre-trial rest best predict the subsequent choice to think about the self. The default network core subsystem classification accuracy is higher than the whole-brain accuracy, suggesting that the whole-brain results are largely driven by the default network core subsystem. We would expect that even though the whole-brain includes the default network core, its accuracy would be lower because of the noise created by other areas of the brain outside of the core system.

**Fig. 3.**
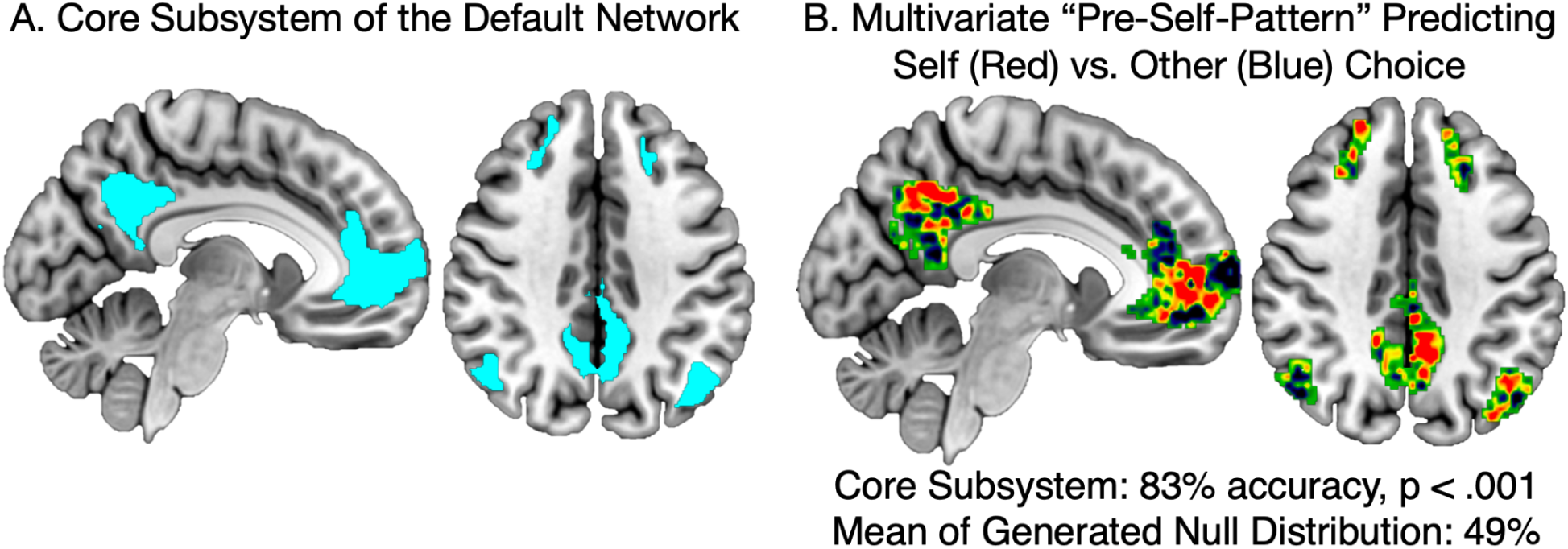
The default network’s core subsystem during pre-trial rest periods predicted the decision to choose to think about the self with 83% accuracy. This accuracy was significantly better (p<.001) than the average classification accuracy in the generated null distribution, which was 49%. (A) depicts the default network’s core subsystem. (B) depicts the multivoxel pattern generated by the SVM to differentiate rest activity that proceeds self-choice rather than other-choice. Red indicates regions where activity predicts self-choice and blue indicates regions where activity predicts other-choice.

We next dug another layer deeper, to understand precisely which default network core regions besides MPFC/BA10 contribute to its classification power. We performed follow-up analyses with all of the ROIs included in Yeo’s default network core subsystem: right middle temporal gyrus, right superior frontal gyrus, left middle frontal gyrus with left superior frontal gyrus, left angular gyrus, right angular gyrus with right middle temporal gyrus, posterior cingulate/precuneus (PCC), and MPFC/BA10. The only classifiers that were significant with an uncorrected threshold were the PCC (69% accuracy, *p* = .02) in addition to MPFC/BA10 (70% accuracy, *p* = .01). To determine if one region was contributing more than the other to the core subystem’s predictive power, we ran an MVPA analysis with the default network core subsystem minus MPFC/BA10 (73% accuracy, *p* = .004), the default network core subsystem minus the PCC (73% accuracy, *p* = .005), the default network core subsystem minus MPFC/BA10 and PCC (59% accuracy, *p* = .17), and MPFC/BA10 and PCC combined (72% accuracy, *p* = .006). We ran statistical tests on a total of 15 classifier models over the course of this analysis. When we correct for multiple comparisons only the default network core subsystem (83% accuracy, *p* = .0001) and the whole brain (77% accuracy, *p* = .001) survive the corrected threshold (*p* < .05/15, or .003). These results in combination with the above suggest MPFC/BA10 and PCC equally contribute to the success of the default network core classifier accuracy, but the activation pattern of the whole default network core subsystem is needed for the best classification power.

### A neural signature that predicts self-focus: Evidence from the resting state scan

#### The Pre-self Pattern in the Default Network’s Core Subsystem during Long Periods of Rest Predict Self-Reported Self-Focus

So far, we have identified a pre-self pattern which predicts self-focused behavior in a forced choice task. Our next goal was to determine if the pre-self pattern generalizes to predict self-focus in another context—a resting state scan. In the beginning of our experiment, participants completed an 8-minute resting state scan and every 2 minutes completed a series of self-reports. They were asked to rate, on a 1-to-5 scale, how much they are thinking about themselves, others, the past, and the future (Fig. 1B). Here, we asked: does the presence of the pre-self pattern preferentially predict self-reported self-focus? We performed an instatement analysis–a within-subjects, TR-to-TR multivariate pattern similarity analysis–assessing the similarity between 1) the pre-self pattern and 2) subjects’ resting state scan pattern in the default network core subsystem. Specifically, each TR of the rest data was masked by the Yeo (2011) default network core subsystem and correlated with the default network core subsystem pre-self pattern (Fig. 4A provides a visualization of the approach). Correlation values were then averaged for the two-minute rest sections and fisher z-transformed. A linear mixed model assessed how self-reported thought content (self, other, future, and past) as well as the section of rest affected the mean correlation. To ensure that the regressors were not introducing collinearity to the model we ran tolerance and variance inflation factors (VIF). Results indicated that collinearity is not an issue in our model: all tolerance values were over 75% (Self = 82, Other = 90, Future = 77, Past = 88, and Section = 93) and all VIF results were well below the threshold of 5 and approaching the baseline of 1 which demonstrates no collinearity (Kutner et al., 2005) (Self = 1.2, Other = 1.1, Future = 1.3, Past = 1.1, and Section = 1.1).

**Fig. 4.**
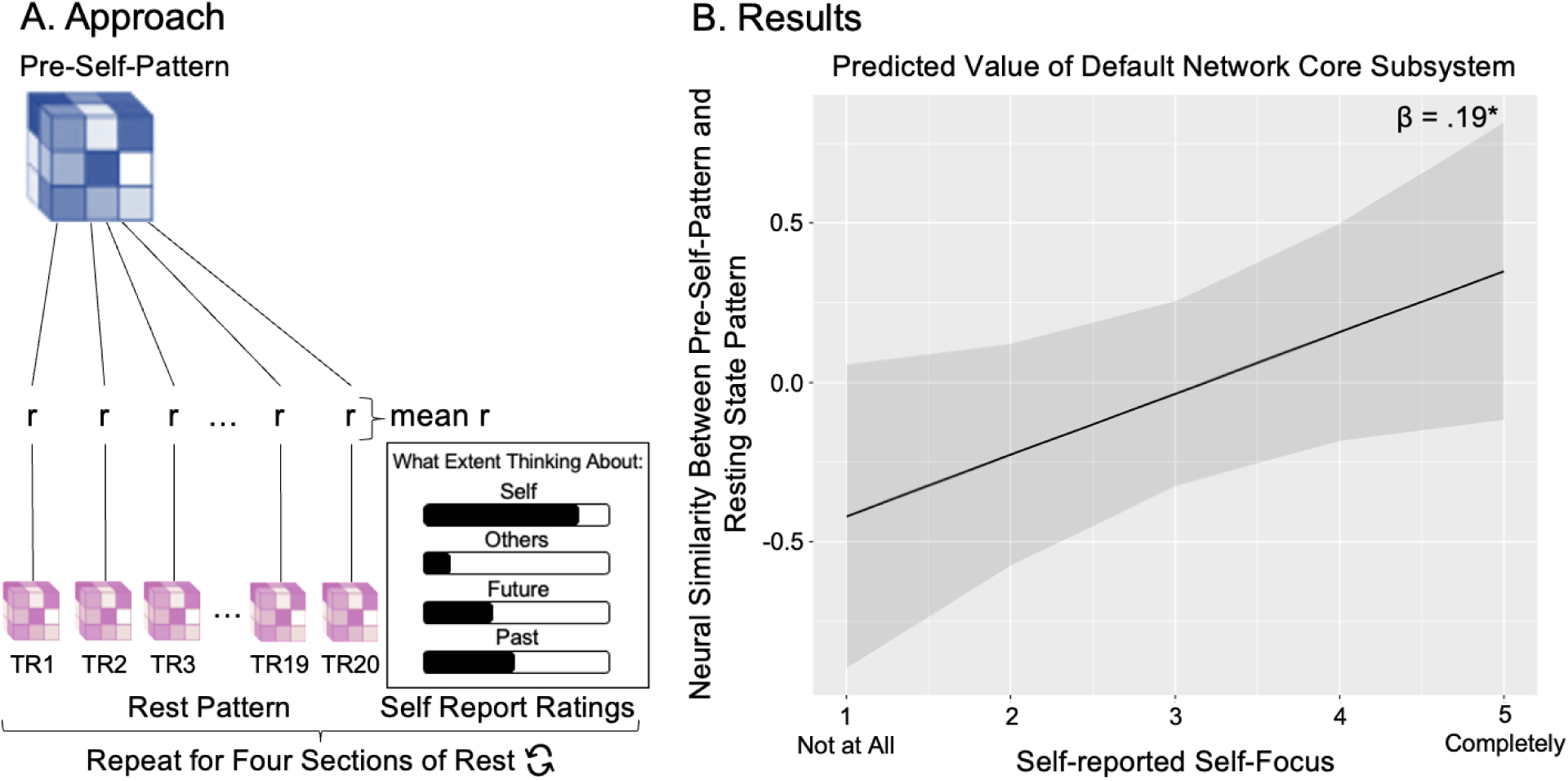
Applying the pre-self pattern to a resting state scan with experience sampling. (A) Our analytic approach started with a within-subjects, TR-to-TR multivariate pattern similarity analysis, assessing the similarity between 1) the pre-self pattern and 2) subjects’ resting state scan pattern in the default network core subsystem. Those correlation values were then averaged for the two-minute rest sections and fisher z-transformed. A linear mixed model assessed how self-reported thought content (self, other, future, and past) as well as the section of rest affected the mean correlation. B) In the results graph, the y-axis displays the normed mean correlation of the resting state neural patterns (in the default network core) with the multivoxel classifying pre-self pattern generated (in the default network core). The x-axis is the self-reported self-focus rating that follows each of the two-minute sections of rest from 1 = “not at all” to 5 = “completely”. The dark gray bars reflect the confidence intervals based on standard error. (+/- 1.96 * SE). We found that the strength of the default network core subsystem multivoxel pattern during rest was significantly related to self-reported self-focus (β = .19, standardized β = 0.09, t(105.1) = 2.03, p = 0.045). In other words, the default network core multivariate pattern derived from short, jittered rest to predict subsequent choice to think about the self, is also able to decode self-focused thought during resting state.

**Fig. 5.**
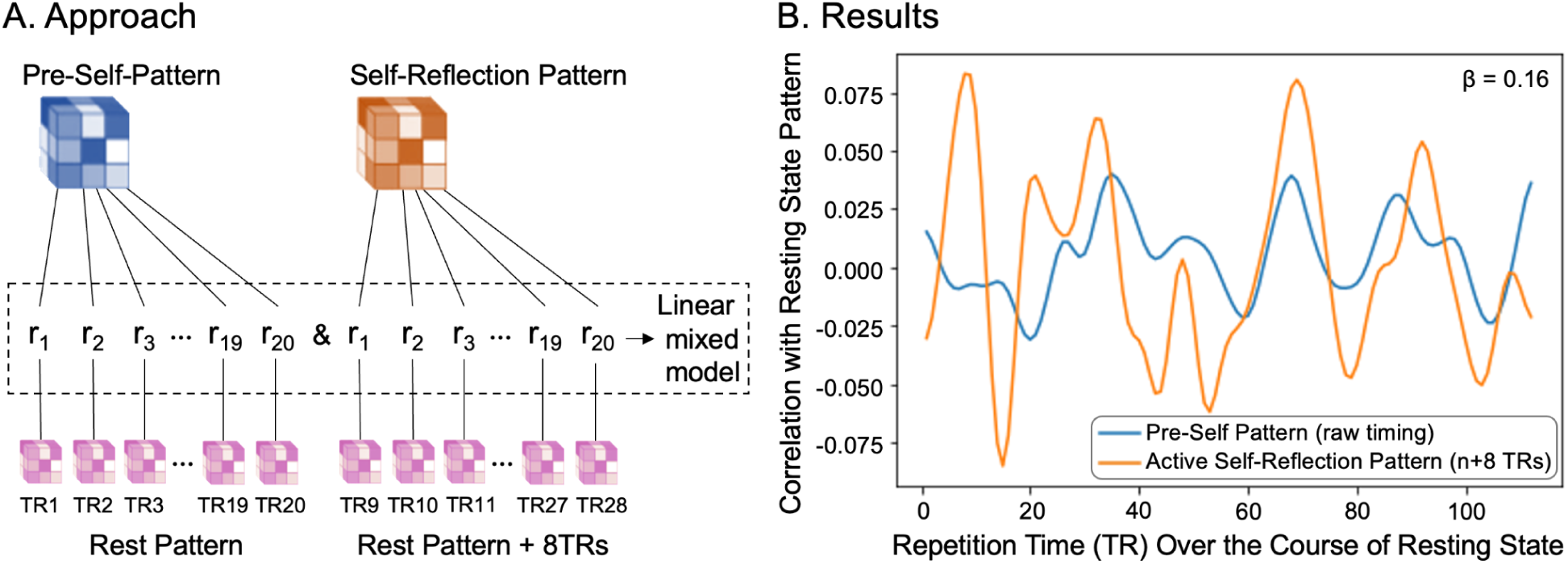
Relationship between Pre-self Pattern and Active Self-Reflection Pattern Over the Resting State Scan (A) depicts the approach using an 8 TR delay: the pre-self pattern (blue) was correlated with each TR (i.e. 1 second) of the resting state scan and the active self-reflection pattern (orange) was also correlated with each TR of the resting state scan. Linear mixed models (Bates et al., 2015) assessed if pre-self pattern correlation strength predicted active self-reflection pattern correlation strength 0-20 TRs later (here 8 TRs later). This allowed us to assess whether the presence of the pre-self pattern temporally predicted the presence of the active self-reflection pattern at multiple lag-times. (B) visualizes the results for a single subject using an 8 TR delay, demonstrating that the presence of the pre-self pattern (blue) predicts the presence of the active self-reflection pattern (orange) 8 seconds later. Note that the active self-reflection pattern strength visualized in orange has been shifted in time (n + 8 seconds) to help visualize its relationship with the pre-self pattern.

A stronger presence of the pre-self pattern corresponded with greater self-reported self-focused thought (β = .19, standardized β = 0.09, *t*(105.1) = 2.03, *p* = 0.045; Fig. 4B) but not other-focused thought (β = 0.07, standardized β = 0.08, *t*(104.1) = .85, *p* = 0.40), future-focused thought (β = -0.01, standardized β = 0.08, *t*(92.6) = -0.10, *p* = 0.92), past-focused thought (β = 0.05, standardized β = 0.07, *t*(86.8) = 0.70, *p* = 0.49) or the section of the resting state scan (β = .06, standardized β = 0.06, *t*(80.1) = 0.91, *p* = 0.37). Follow-up analyses with the whole brain, dmPFC and MTL subsystems, as well as MPFC/BA10 and PCC examined individually, did not produce significant results (β’s < 0.14, *p’s* > 0.10). Thus, multivariate patterns in the default network core subsystem as a whole—derived from short, jittered rest to predict subsequent choice to think about the self—is also able to predict self-reported, self-focused thought during a resting state scan.

#### The Pre-self Pattern Temporally Predicts the Active Self-Reflection Neural Pattern during a Resting State Scan

We have shown the pre-self pattern predicts behavioral markers of self-focus: self-focused decisions and self-reported self-focus. Could the pre-self pattern even predict a marker of self-focus that participants do not necessarily report on? To answer this question, we computed, for each subject, a multivariate pattern that reflected their active self-reflection in our final fMRI task, in which they answered questions about themselves such as “am I an ambitious student?” (Fig. 1C). We then used instatement analysis to test whether the presence of the pre-self pattern precedes the presence of this active self-reflection pattern in the core default network subsystem. We restricted our analyses to the core subsystem because of the accumulating evidence so far that distributed patterns in this network meaningfully predict self-focus.

We performed an instatement analysis, taking each participant’s multivariate self-reflection pattern in the core subsystem from the final fMRI task and assessing its neural pattern similarity with each second (i.e., TR) of the resting state scan. We then assessed if resting state neural pattern similarity to the pre-self pattern temporally predicted neural pattern similarity to the self-reflection-pattern at different timing delays. Specifically, linear mixed models (Bates et al., 2015) assessed if pre-self pattern correlation strength predicted active self-reflection pattern correlation strength 0-20 TRs later (See Fig. 5A for a visualization of the approach). We chose to use this data driven approach with multiple TR lag-times to avoid arbitrarily selecting a time delay as well as to better understand the dynamic temporal relationship between the presence of the “pre-self” pattern and active self-reflection pattern. Analyses are performed on fisher z-transformed correlation values.

It is noteworthy that the pre-self pattern and each subjects’ self-reflection pattern demonstrated small correlations (-.03 < *r*’s < .08). We therefore ran the linear mixed models with 0-20 TR delays: 1) with the strength (i.e., correlation) of the pre-self pattern predicting the strength of the active self-reflection-pattern X TRs later and 2) with the strength (i.e., correlation) of the active self-reflection pattern predicting the strength of the pre-self pattern X TRs later. To account for the number of statistical tests performed (41), we used a bonferroni-corrected significance value of *p* < 0.00122 (i.e., .05×41).

Our first observation further indicated that the pre-self pattern and active self-reflection patterns are not redundant with one another, and instead are distinct brain states. The presence of the pre-self pattern and the presence of an active self-reflection pattern were *negatively* correlated when we used no TR delay (β = -0.16, standardized β = 0.03, *t*(15340) = -4.70, *p* <0.00001) as well as when there was a 1 or 2 TR delay of the pre-self pattern predicting the active self-reflection pattern (1TR: β = -2.47, standardized β = .04, *t*(15210) = -7.16, *p* <0.00001; 2TR: β = -1.27, standardized β = 0.04, *t*(15080) = -3.65, *p* = 0.0003) These results demonstrate that our pre-self pattern is capturing neural activity that is distinct and different from active self-reflection. The negative correlations suggest that when someone is more strongly in the pre-self pattern “state” they are less likely to be in the active self-reflection “state”, pointing to their distinctness.

Our second observation was that there is a temporal relationship between the pre-self pattern and active self-reflection pattern. The strength of the pre-self pattern significantly predicted the strength of the active self-reflection pattern 8 TRs later (β = 0.16, standardized β = 0.04, *t*(14310) = 4.55, *p* < 0.00001). Fig. 5B depicts this finding for one example participant. This temporal relationship also appeared 11 TRs later (β = 0.16, standardized β = 0.04, *t*(13930) = 4.44, *p* = 0.00001), 13 TRs later (β = 0.14, standardized β = 0.04, *t*(13670) = 3.94, *p* = 0.00008), and 14 TRs later (β = .15, standardized β = 0.04, *t*(13540) = 4.08, *p* = 0.00005). In contrast, the active self-reflection patterns did not significantly predict a stronger presence of the pre-self pattern with any time delay (*t*’s -2.37-2.55, *p*’s 0.01-0.96), apart for the 1 and 2 TR delays showing a negative relationship that are reported in the above paragraph. See Supplementary Table 1 for a full list of the results for each TR lag. These results suggest multivariate patterns in the default network core subsystem—derived from short, jittered rest to predict subsequent choices to think about the self—are also able to temporally predict neural indices of active self-reflection during a resting state scan, even when the neural indices of active self-reflection are clearly distinct from the pre-self pattern by virtue of them being negatively correlated at the same moment in time.

We also capitalized on the active self-reflection pattern to test whether the relationship between the pre-self pattern and self-reported self-focus (reported in the section above) is independent of a relationship between the active self-reflection pattern and self-reported self-focus. A linear mixed model, that included both the pre-self pattern and the active self-reflection pattern as independent predictors of self-reported self-focus showed that there was still a significant relationship between pre-self pattern instatement and self-reported self-focus (β = .20, standardized β = 0.10, *t*(102.4) = 2.08, *p* = 0.041). Indeed, when we looked at variance inflation factors, VIF results were well below the threshold of 5 and approaching the baseline of 1 which demonstrates no collinearity (Kutner et al., 2005) (‘pre-self’ pattern = 1.01 and self-reflection pattern = 1.02). This further suggests that the pre-self pattern is not redundant with active self-reflection.

## Study 2: Implications for Internalizing and Out-of-Sample Testing

The results from Study 1 suggest that we can decode the bias towards self-focus in our own dataset. The pre-self pattern predicts self-focused decisions, subjective self-focus during rest, and the presence of active self-reflection neural patterns a few seconds later. Self-focused thought is implicated in mental health conditions, particularly internalizing disorders such as anxiety and depression (Baez et al., 2023; du Pont et al., 2018; Watkins, 2008; Watkins & Roberts, 2020). Could the pre-self pattern be used to predict internalizing scores in an entirely separate sample of participants? If so, this would have great utility: it would suggest we derived a neural marker that, down the line, could be used to identify a person’s vulnerability to internalizing conditions. In a first step towards this goal, we took the pre-self pattern, created in our 32 subject dataset, and applied it to a larger dataset to show that the pattern translates not just across tasks but also across datasets and subjects.

## Methods

### Human Connectome Project Participants

We applied our pre-self pattern to data from the first resting state scan of the Human Connectome Project, hereafter referred to as the Human Connectome Project dataset (Van Essen et al., 2013). The dataset is openly accessible and consists of a large sample of neurotypical individuals. Data from the Human Connectome Project are publicly available in the online Human Connectome Project repository (https://db.humanconnectome.org/; fMRI data are in the subfolders rfMRI_REST1_RL, behavioral data are in the Restricted Data file). We utilized neural and behavioral data from one thousand eighty-six individuals (age 22–37 years, mean age 28.8; 588 female and 498 male; 817=White, 63=Asian/Nat. Hawaiian/Othr Pacific Is., 158=Black or African Am., 2=Am. Indian/Alaskan Nat., 28=More than one, 18=Unknown or Not Reported; 979=Not Hispanic/Latino, 94=Hispanic/Latino, 13=Unknown or Not Reported).

### fMRI Collection

The resting state scan that we used in our analysis was 14 minutes 33 seconds long and was the first functional scan done on participants’ first day in the lab. The fMRI data were acquired using a 3T Skyra scanner with 2 mm isotropic voxels and a TR of 0.72 s (for more acquisition details see Van Essen et al., 2013). Each run comprised 1200 scan volumes, and there was a single run for each participant. We used minimally preprocessed voxelwise fMRI data. See Glasser et al. (2013) for further details about data preprocessing.

### Residual Image Calculation

Consistent with Study 1’s data analysis pipeline and prior resting state research (Meyer et al., 2019; Tambini & Davachi, 2019), we also calculated residual images for the Human Connectome Project resting state scan. Residual images were calculated with a general linear model that included nuisance regressors for the six motion parameters (x, y, and z directions and roll, pitch, and yaw rotations), each motion parameter’s derivative and square of the derivative, and linear drift. We additionally regressed out TRs in nonsteady state and TRs that exhibited spikes of motion found from global signal outliers and outliers derived from frame differencing (each 3 SDs). Finally, we used a smoothing kernel of 6. All analyses of the Human Connectome Project resting state scan were run on the residual images saved from those models.

### Linking Pre-self Pattern Instatement and Internalizing In the Human Connectome Project Dataset

For each Human Connectome participant, we first carried out an instatement analysis in which we performed a TR-to-TR pattern matching analysis with our pre-self pattern–the classifier pattern generated by the multivoxel pattern analysis of the default network core subsystem. Each TR of the 14.5 minutes of rest was masked by the Yeo (2011) default network core subsystem ROI and then correlated with the default network core subsystem pre-self pattern. This created a timecourse for each subject where each data point indicated the extent to which the participant’s neural activity matched our pre-self pattern at that particular time point during rest (Fig. 6A).

**Fig. 6.**
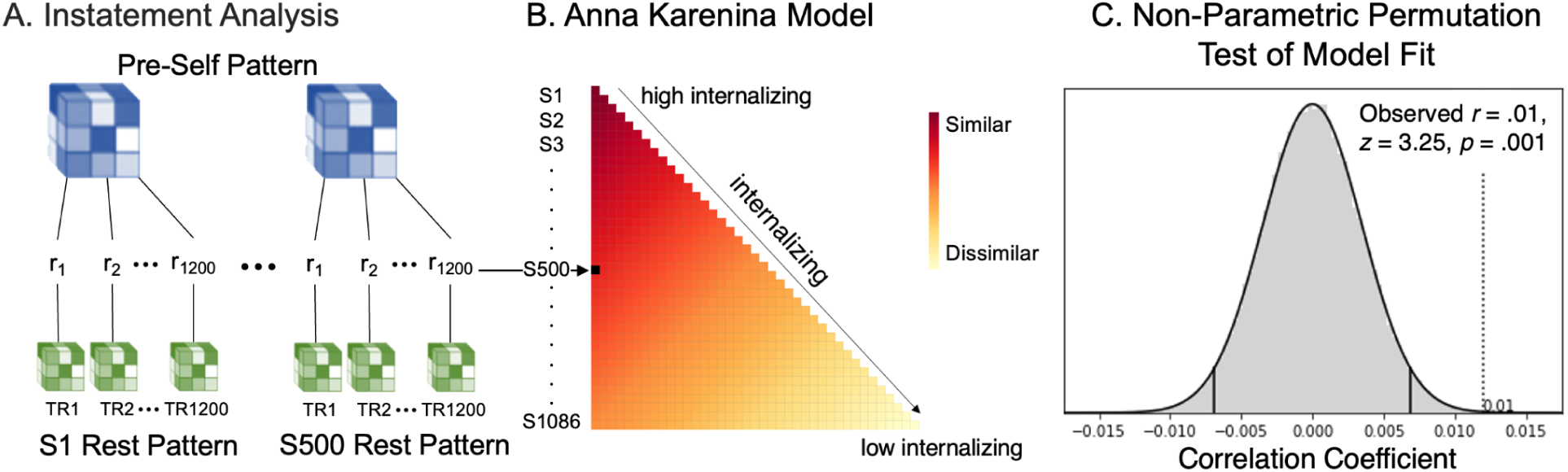
Intersubject Representation Similarity Analysis (IS-RSA) with Anna Karenina Model: High Internalizations have a Similar Pre-Self Pattern Timecourse during a Resting State Scan The Anna Karenina model testing whether high internalizers move in and out of the preself pattern similarly over the course of a resting state scan. (A) Similarity in the timing of the pre-self pattern strength was computed for all pairs of Human Connectome Project participants and compared to (B) the theoretical model that high internalizers show a similar timecourse to one another while low internalizers show idiosyncratic timecourses. (C) A non-parametric, Mantel permutation test (see Methods) showed the internalizing Anna Karinina model was significant.

#### Average Pre-self Pattern Instatement During Rest and Internalizing

We arbitrated between two ways in which instating the pre-self pattern may relate to internalizing. The first possibility is that an overall greater degree of pre-self pattern instatement predicts internalizing, given that individuals higher on this trait dimension report an overall greater amount of self-focus. To test the first possibility, after we performed the instatement analysis detailed above, we averaged pre-self pattern instatement values across all TRs for each participant. We then created a linear model in R with the *stats* package (R Core Team, 2022) to determine if participant’s internalizing scores significantly predicted their average pre-self pattern instatement during rest.

#### Intersubject Representational Similarity Analysis

The second possibility is that individuals who score high on internalizing show a similar timecourse to the presence of their pre-self pattern, moving in and out of the pre-self pattern similarly over the course of the resting state scan. To test this prediction explicitly, we used an intersubject representational similarity analysis (IS-RSA; Finn et al., 2020) to test an “Anna Karenina” model. Statistically, Anna Karenina models test if “all high (or low) scorers are alike; each low (or high) scorer is different in his or her own way,” with the researcher specifying whether they predict similarity among high or low scorers (Chang et al., 2020; Finn et al., 2020). In our case, we specified the model to test if all high internalizers move in and out of the pre-self pattern alike, while low internalizers move in and out of the preself pattern in their own unique ways.

Following suggestions from Finn et al. (2020), we modeled subject similarity based on participants’ absolute position on the internalizing scale by computing the mean internalizing rank (i.e., (rank(i) + rank(j))/2). This approach is the appropriate choice for Anna Karenina models, as opposed to Euclidean distance or other relative distance measures (which are used in nearest neighbor models), because Euclidean distance between subjects would be agnostic to where on the internalizing continuum participants fall. We hypothesized that high-internalizing individuals exhibit greater similarity among themselves, while low-internalizing individuals are less similar to both high-internalizing individuals and low-internalizing individuals. Therefore, we modeled subject similarity via mean internalizing rank, which captures this relationship. See Fig. 6B for a depiction of the Anna Karenina model.

IS-RSA requires computing two subject-by-subject matrices (here, one matrix for subjects’ internalizing scores and one matrix for subjects’ pre-self pattern instatement timecourses) and statistically comparing them to one another. To create the internalizing score matrix, subjects’ internalizing scores were turned into ranks, making low internalizing subjects ranked low and high internalizing subjects ranked high (range of ranks = 0-1085 for N=1086). We then computed the mean of these ranks for every subject pair to generate our 1086*1086 intersubject mean internalizing matrix. Next, we Pearson correlated each subject’s pre-self pattern instatement timecourse with every other subjects’ pre-self pattern instatement timecourse to populate the neural representational similarity matrix. This instatement timecourse similarity matrix was organized as a function of participant’s internalizing scores, so that participants higher on internalizing appeared at the top and those lower on internalizing appeared on the bottom (Fig 6A and B).

Next, to test our hypothesized link between high internalizing and similar pre-self pattern timecourses during rest, we spearman correlated our intersubject internalizing and instatement timecourse similarity matrices (specifically, correlating only the lower triangles of these symmetric matrices). We used spearman correlations, as is the protocol in the Representational Similarity Analysis literature (Kriegeskorte et al., 2008), because the increase in pre-self pattern instatement timecourse similarity may not be linear to either the increase in the internalizing of a subject-pair or the increase in the internalizing similarity of a subject-pair.

To test our model’s statistical significance, we needed to account for each subject appearing in the model multiple times—as we compared every subject to every other subject indicating each subject appears N-1 (1085) times in our model. To account for this non-independence in our data, we tested our model with a non-parametric, Mantel permutation test, as has been done in previous research (Finn et al., 2018; Iyer et al., 2024; Sava-Segal et al., 2023). Specifically, we randomly shuffled the identity associated with subjects’ instatement timecourses (e.g. each subject’s (intact) instatement timecourse was relabeled with a different subject’s identity) 100,000 times, each time correlating the resulting simulated instatement timecourse similarity matrix with our unshuffled Anna Karenina calculated internalizing matrix. This created a null distribution of IS-RSA correlation values. We then quantified the probability that our results were produced by chance by computing the proportion of times our simulated null correlation value exceeded our observed model-data correlation. Finally, to determine statistical significance we compared this probability to a significance threshold of alpha = .05. As a follow-up, we also performed IS-RSA Anna Karenina analyses separately for each of the three subscales within the internalizing measure–(1) Anxiety and Depression, (2) Withdrawal, and (3) Somatic Complaints. We replicated our previous methodology three times, substituting each subscale measure for the overall internalizing measure.

## Results

### A neural signature that predicts self-focus: Evidence from the Human Connectome Project

#### In the Human Connectome Project Dataset, High Internalizers Move In and Out of the Pre-self Pattern Similarly Over Time

We arbitrated between two ways in which instating the pre-self pattern during rest may relate to internalizing. The first possibility is that an overall greater degree of pre-self pattern instatement predicts internalizing, given that individuals higher on this trait dimension report an overall greater amount of self-focus. To test the first possibility, for each participant, we computed the average strength (i.e. correlation) in the core subsystem between the pre-self pattern and their rest pattern across all TRs of the resting state scan. We then tested if internalizing scores predicted the average amount of pre-self pattern instatement during rest. Internalizing scores were not significantly related to the overall amount of pre-self pattern instatement (β = -0.00001, standardized β = -0.02, *t*(1085) = -0.62, *p* = 0.54). Put simply, high internalizers do not instate the pre-self pattern to a greater degree than low internalizers.

Next, we tested the second possibility: individuals higher on the internalizing trait dimension move in and out of the pre-self pattern similarly over the course of rest. This possibility is consistent with recent research showing that individuals with similar trait characteristics show similar timecourses of neural responding (Finn et al., 2018; van Baar et al., 2021), including over the course of resting state scans (Iyer et al., 2024; Liu et al., 2019; Qiu et al., 2024). To test this possibility, we performed an Anna Karenina intersubject representational similarity analysis (IS-RSA), which can test whether individuals who score high on a trait dimension show similar neural responses, whereas individuals who score low on the same trait dimension are neither similar to other low nor high scorers (i.e., there is no systematic structure to low scorers’ neural responding; Fig. 6). We chose this model because we did not expect any particular timing with respect to when low internalizers instate their pre-self pattern. In contrast, we reasoned high internalizers may instate their pre-self pattern in systematic ways (and hence similarly) over time. The Anna Karenina model was significant (*r* = 0.01, *z* = 3.25, *p* = 0.001, Mantel permutation test; Fig. 6C). In other words, people high on internalizing, a clinical variable highly related to maladaptive self-focus, move in and out of the pre-self pattern similarly over time throughout a resting state scan. For visualization purposes, Fig. 7 shows pre-self pattern instatement timecourses from four Human Connectome participants: two who score high (left panel) and two who score low (right panel) on internalizing.

**Fig. 7.**
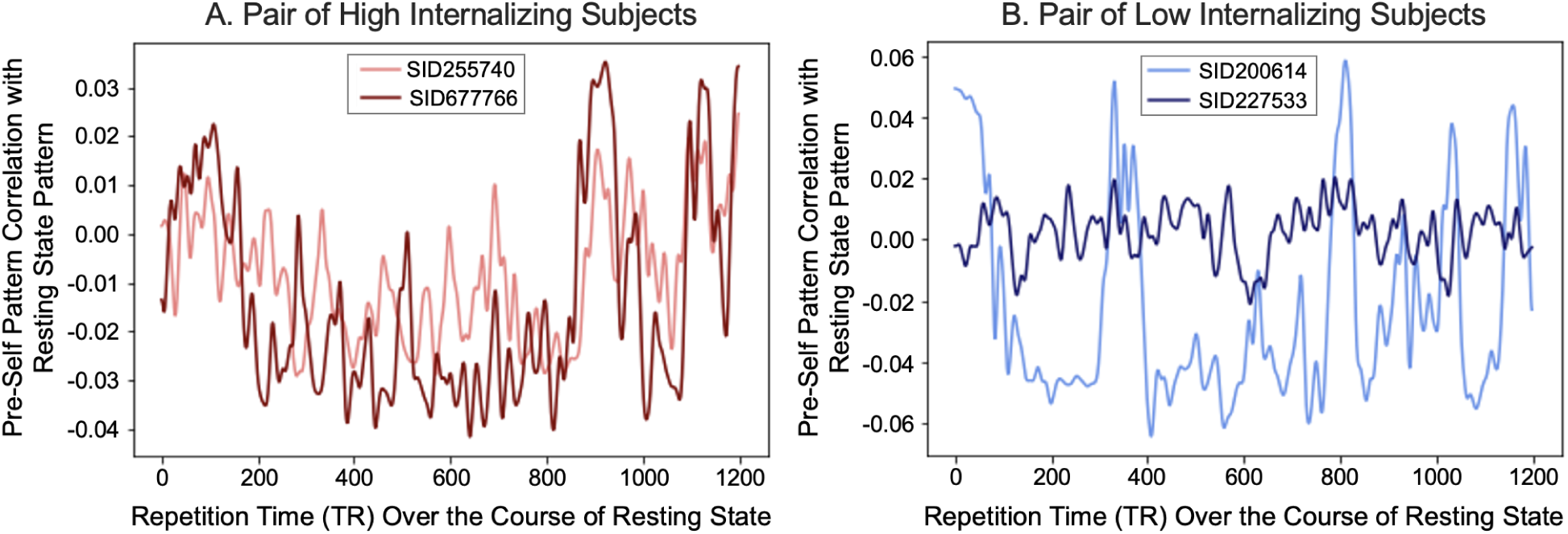
Intersubject Representational Similarity Analysis (IS-RSA): Example Subjects’ Pre-Self Pattern Instatement Timecourse For our IS-RSA we compared the timecourse of individuals’ pre-self pattern instatement strength with an Anna Kerenina model of internalizing such that the higher a pair’s internalizing rank, the higher their similarity in pre-self pattern instatement timing, whereas the lower their internalizing rank, the more idiosyncratic their timing. A. shows two high internalizing subjects (90th percentile) pre-self pattern instatement timecourse while B. shows two low internalizing subjects (10th percentile) timecourse.

The internalizing measure used in the Human Connectome Project includes three subscales: (1) anxiety and depression, (2) withdrawal, and (3) somatic symptoms. Importantly, the first two dimensions relate to psychological self-focus more so than a focus on one’s bodily sensation. As a follow-up, we performed IS-RSA analyses, separately for each of three of the subscales within the internalizing measure. Both the Anxiety and Depression subscale and the Withdrawal subscale showed a significant Anna Karenina effect (Anxiety and Depression: *r* = 0.014, *z =* 4.79*, p* < 0.00001; Withdrawal: *r* = .01, *z =* 2.86*, p* = 0.004). Somatic Complaints, however, did not show a significant effect (*r* = 0.0001, *z =* 0.02*, p* = 0.98).

Supplementary materials detail follow-up analyses that further confirm that high internalizers similarly instate the pre-self pattern over time during rest. Briefly here, the results could not be attributed to 1) similar pre-self pattern instatement timecourses for similar levels of internalizing scores across the sample, rather than *high* internalizing scores specifically or 2) high internalizers showing different degrees of variability in their pre-self pattern instatement during rest. Collectively, the Human Connectome Project findings suggest that people high on internalizing, particularly anxiety, depression, and withdrawal, show similar timing to the presence of the pre-self pattern during a resting state scan, suggesting there may be a systematic, temporal structure to their spontaneous, self-focused thought.

## Discussion

Although self-focus is a ubiquitous part of human psychology, the underlying processes that generate interest in ourselves remain to be fully characterized. We identified a brain state that when entered during brief mental breaks predicts participants’ immediately following decision to focus on themselves. In Study 1, we found a multivariate neural pattern in the default network’s core subsystem, which includes MPFC/BA10, that can decode the subsequent decision to focus on the self with high accuracy. When we apply this ‘pre-self pattern’ to other outcomes in our dataset, its presence in the default network’s core subsystem also predicts self-reported self-focus during a resting state scan. It is even capable of temporally predicting the presence of a neural pattern capturing active self-reflection during a resting state scan. In Study 2, we applied the pre-self pattern to data from the Human Connectome Project and found our pre-self pattern significantly related to internalizing scores in this different sample of participants. Individuals with high internalizing scores moved into and out of the pre-self pattern in similar ways during a resting state scan, whereas individuals with low internalizing scores showed no similar timing to their pre-self pattern. This result offers a unique way to think about how internalizing relates to self-focus–there may be a systematic timing to high internalizers’ spontaneous, self-focused thought in the core default network.

What is going on–in terms of a psychological process–in participants’ brains when we observe the pre-self pattern? Importantly, while the pre-self pattern is modestly correlated with a pattern reflecting active self-reflection (from a task in which participants were instructed to reflect on themselves), it is not redundant with the active self-reflection pattern. Rather, in any given moment, the presence of the pre-self pattern is negatively correlated with the presence of the active self-reflection pattern, yet it can temporally predict the later presence of the active self-reflection pattern. The pre-self pattern and active-self reflection patterns also explain unique variance in participants’ self-reported self-focus. All together, the data suggests whatever psychological process may be captured by our pre-self pattern, it appears distinct from active self-reflection.

There may be at least three possible psychological phenomena captured by our pre-self pattern. First, the pre-self pattern may signify a “loading” of the self; a coming online of a self-schema through which to view and interact with the world. Consistent with this view, some posit that we constantly exist in a framework of the self, as we understand our world through our own experiences and embodied sense of self, which we constantly exist in (Christoff et al., 2011). The second possibility is related to the first: the pre-self pattern could reflect the persistent cognitive accessibility of the self (Bruner, 1957; den Daas et al., 2013; Libet et al., 1983). Cognitive category accessibility was first proposed by Bruner (Bruner, 1957) and suggests that the attentional framework we are in directly influences what we subsequently perceive or think about. Thus, the pre-self pattern may reflect an attentional state that facilitates quick access to self-knowledge. Third, it is possible the pre-self pattern represents a motivational state. Specifically, participants may be experiencing a temptation or impulse to think about the self. This may be an urge that they could potentially resist or succumb to, perhaps with significant implications for mental health, specifically rumination. Future research will help arbitrate between these competing possibilities. Though we cannot be certain what psychological phenomena the pre-self pattern is capturing, its observation is consistent with predictive coding accounts of brain function, which broadly suggest endogenous, default brain states predict subsequent perception and cognition (Friston, 2018; Y. Huang & Rao, 2011; Hutchinson & Barrett, 2019).

Another important question to address about the pre-self pattern is whether it captures conscious preparation or deliberation to think about the self versus a more implicit process. Prior work has observed that nonconscious pre-stimulus activity can predict subsequent decisions (Huang et al., 2014; Soon et al., 2008, 2013). Relevantly, previous fMRI and intracranial recording research has found that neural activity in MPFC and PCC–the two regions in our data with the strongest classification power to predict subsequent self-focused thought–can predict subsequent decisions to press a button or perform a math task 2-7 seconds before the participant reports making a conscious choice (Fried et al., 2011; Soon et al., 2008, 2013). This research is related to previous EEG research into the “readiness potential”, a build-up of activity in the prefrontal cortex prior to the conscious decision to move (Libet et al., 1983; Schurger et al., 2021). Researchers theorize that when this spontaneous neural activity passes a threshold or coincides with a signal from a different brain region, it will nudge a participant into motion without prior, conscious deliberation (Murakami et al., 2017; Schurger et al., 2012, 2021). This fits with our current findings, as our self-vs. other-focus decision task allows us to capture a (perhaps preconscious) build-up of neural activity before the choice to be self-focused. We then show how spontaneous fluctuations of the preconscious self-focus activity (e.g. our pre-self pattern instatement) is tied with subsequent conscious thought (e.g. self-reported self-focus and the neural pattern of active self-reflection).

Our results offer a new perspective on psychological theories of the self. Social psychological theories argue humans are interested in themselves to meet both an epistemic goal to understand their reality and a belonging goal to feel connected to groups of people (Kwang & Swann, 2010; Leary, 2007). Our results offer a new twist on these ideas – there is a momentary brain state we enter reflexively, even in just a few second mental break, that may help ensure we default towards trying to meet these goals. Future work can test this possibility more directly, for example by examining 1) whether engaging the pre-self pattern during brief mental breaks guides decisions that help meet epistemic and belonging goals and 2) whether manipulating the need to achieve these goals increases instatement of the pre-self pattern during brief mental breaks.

The pre-self pattern may also update understanding of cognitive phenomena related to the self. For instance, the “self-reference effect” refers to the robust finding that people are more likely to remember information related to themselves (vs. unrelated to themselves). Historically, this area of research suggests the self-reference effect occurs because people have highly elaborate and organized self-schemas, which facilitates higher quality processing of self-relevant content when encoding new information (Symons & Johnson, 1997). Our work suggests another, complementary possibility – because the pre-self pattern emerges by default during momentary mental breaks, having it active just prior to encoding new stimuli may make stimuli that is related to the self more efficiently incorporated into self-knowledge. This idea fits with the concept of “preplay” from animal research. This work shows that the neurons that are already active in a rodent during brief rest before completing a novel spatial maze guide behavior and subsequent memory of their maze trajectory (Buhry et al., 2011). Future work can test whether spontaneously instating the pre-self pattern during brief mental breaks facilitates new learning of self-related information moments later.

The pre-self pattern is also useful because it predicts self-focus in multiple ways and even across different datasets. This means that it could be utilized as a decoding tool in future research on the role of self-focus in other relevant behaviors. For example, self-focus during conversation corresponds with worse relationship quality (Schwartz-Mette & Rose, 2009) and reducing self-concept accessibility has been suggested to help us engage with other people’s points-of-view (Kaufman & Libby, 2012). Yet, objective markers of these phenomena, as well as a clear explanation as to how and why they occur, remains to be determined. The pre-self pattern provides a neural signature to better understand the bias towards self-focus in multiple aspects of everyday life.

The pre-self pattern may also be a useful decoding or investigative tool in mental health research. Among Human Connectome Project participants, similarly moving in and out of the pre-self pattern during a resting state scan corresponded with internalizing symptoms, particularly depression, anxiety, and withdrawal. Rumination–repetitive and recurrent negative thinking about oneself (Watkins, 2008)–is thought to play a key role in internalizing disorders (Baez et al., 2023; du Pont et al., 2018; Watkins & Roberts, 2020). Yet, the neural mechanism that explains why and how rumination spontaneously occurs is not fully understood, although past work has associated the default network with depression and anxiety (Coutinho et al., 2016; Sheline et al., 2009). The pre-self pattern may offer key insight into the basic mechanisms underlying rumination. The tendency to systematically move in and out of the pre-self pattern may bias high internalizers towards the repetitive, negative thoughts characteristic of rumination. Moreover, the pre-self pattern may be applied to patient populations and/or individuals at risk for developing internalizing disorders to help predict, and perhaps eventually offset, maladaptive self-focus.

Self-focus is a pervasive human phenomenon that plays a vital role in our lives—from meeting our personal needs to understanding our social standing. However, in its most pernicious forms, self-focus is a risk and maintenance factor for internalizing disorders such as depression and anxiety (Ingram, 1990; Just & Alloy, 1997; Kuehner & Weber, 1999; Nolen-Hoeksema, 1991, 2000; Nolen-Hoeksema et al., 1994). This paper documents a neural signature–the pre-self pattern in the core subsystem of the default network–that biases people towards self-focus. The pre-self pattern predicts self-focused decisions, subjective self-focus during rest, and the presence of active self-reflection neural patterns a few seconds later. Moreover, the timing of the pre-self pattern in the core subsystem during a resting state scan predicted internalizing scores in a large sample of participants from the Human Connectome Project. With this pre-self pattern in hand, we are one step closer towards understanding why humans cannot help but focus on themselves, as well as how this process goes awry in mental health conditions.

## Acknowledgements and Funding

This work was supported by an R01 grant from NIMH awarded to Dr. Meghan L. Meyer. Courtney Jimenez, Sasha Brietzke, Christina Huber, and Megan Hillis assisted with data collection. Luke Chang provided valuable advice on data analysis decisions.

## Author Contributions

D.G. and M.M. designed research; D.G. performed research; D.G. analyzed data; and D.G. and M.M. wrote the paper.

## Competing Interests

The authors declare no competing interest.

## Data Availability

Data from our experiment can be found here: https://osf.io/8gefx/: and Human Connectome Data that we analyzed can be found here: https://db.humanconnectome.org/

## Notes

https://osf.io/8gefx/

### Competing Interest Statement

The authors have declared no competing interest.

### Summary of Updates

Major updates to the HCP analysis and to the paper structure.

## References

1. Addante, R. J., de Chastelaine, M., & Rugg, M. D. (2015). Pre-stimulus neural activity predicts successful encoding of inter-item associations. NeuroImage, 105, 21–31.

2. Aly, M., & Turk-Browne, N. B. (2016). Attention promotes episodic encoding by stabilizing hippocampal representations. Proceedings of the National Academy of Sciences of the United States of America, 113(4), E420–E429.

3. Andrews-Hanna, J. R., Kaiser, R. H., Turner, A. E. J., Reineberg, A. E., Godinez, D., Dimidjian, S., & Banich, M. T. (2013). A penny for your thoughts: dimensions of self-generated thought content and relationships with individual differences in emotional wellbeing. Frontiers in Psychology, 4, 900.

4. Andrews-Hanna, J. R., Reidler, J. S., Huang, C., & Buckner, R. L. (2010). Evidence for the Default Network’s Role in Spontaneous Cognition. Journal of Neurophysiology, 104(1), 322–335.

5. Baez, L. M., Puccetti, N. A., Stamatis, C. A., Jaso, B. A., Timpano, K. R., & Heller, A. S. (2023). Identifying real-world affective correlates of cognitive risk factors for internalizing disorders. Emotion, 23(3), 678–687.

6. Bates, D., Mächler, M., Bolker, B., & Walker, S. (2015). Fitting Linear Mixed-Effects Models Using lme4. Journal of Statistical Software, 67, 1–48.

7. Berman, M. G., Peltier, S., Nee, D. E., Kross, E., Deldin, P. J., & Jonides, J. (2011). Depression, rumination and the default network. Social Cognitive and Affective Neuroscience, 6(5), 548–555.

8. Bischoping, K. (1993). Gender differences in conversation topics, 1922–1990. Sex Roles, 28(1), 1–18.

9. Bressler, S. L., & Menon, V. (2010). Large-scale brain networks in cognition: emerging methods and principles. Trends in Cognitive Sciences, 14(6), 277–290.

10. Bruner, J. S. (1957). On perceptual readiness. Psychological Review, 64(2), 123–152.

11. Buckner, R. L., & DiNicola, L. M. (10/2019). The brain’s default network: updated anatomy, physiology and evolving insights. Nature Reviews. Neuroscience, 20(10), 593–608.

12. Buhry, L., Azizi, A. H., & Cheng, S. (2011). Reactivation, replay, and preplay: how it might all fit together. Neural Plasticity, 2011, 203462.

13. Chang, L. J., Gianaros, P. J., Manuck, S. B., Krishnan, A., & Wager, T. D. (2015). A sensitive and specific neural signature for picture-induced negative affect. PLoS Biology, 13(6), e1002180.

14. Chang, L., Jolly, E., Cheong, J. H., Burnashev, A., Chen, A., Clark, M., Frey, S., & Fitzpatrick, P. (2023). *cosanlab/nltools* (Version 0.5.0). 10.5281/zenodo.10059041

15. Chang, L., Manning, J., Baldassano, C., de la Vega, A., Fleetwood, G., Geerligs, L., Haxby, J., Lahnakoski, J., Parkinson, C., Shappell, H., Shim, W. M., Wager, T., Yarkoni, T., Yeshurun, Y., & Finn, E. (2020). naturalistic-data-analysis/naturalistic_data_analysis: Version 1.0. 10.5281/zenodo.3937849

16. Christoff, K., Cosmelli, D., Legrand, D., & Thompson, E. (2011). Specifying the self for cognitive neuroscience. Trends in Cognitive Sciences, 15(3), 104–112.

17. Cohen, J. R., Asarnow, R. F., Sabb, F. W., Bilder, R. M., Bookheimer, S. Y., Knowlton, B. J., & Poldrack, R. A. (2010). Decoding developmental differences and individual variability in response inhibition through predictive analyses across individuals. Frontiers in Human Neuroscience, 4, 47.

18. Coutinho, J. F., Fernandesl, S. V., Soares, J. M., Maia, L., Gonçalves, Ó. F., & Sampaio, A. (2016). Default mode network dissociation in depressive and anxiety states. Brain Imaging and Behavior, 10(1), 147–157.

19. Craik, F. I. M., Moroz, T. M., Moscovitch, M., Stuss, D. T., Winocur, G., Tulving, E., & Kapur, S. (1999). In Search of the Self: A Positron Emission Tomography Study. Psychological Science, 10(1), 26–34.

20. den Daas, C., Häfner, M., & de Wit, J. (2013). Sizing opportunity: Biases in estimates of goal-relevant objects depend on goal congruence. Social Psychological and Personality Science, 4(3), 362–368.

21. Denny, B. T., Kober, H., Wager, T. D., & Ochsner, K. N. (2012). A Meta-analysis of Functional Neuroimaging Studies of Self- and Other Judgments Reveals a Spatial Gradient for Mentalizing in Medial Prefrontal Cortex. Journal of Cognitive Neuroscience, 24(8), 1742–1752.

22. Dunbar, R. I. M., Marriott, A., & Duncan, N. D. C. (1997). Human conversational behavior. Human Nature, 8(3), 231–246.

23. du Pont, A., Rhee, S. H., Corley, R. P., Hewitt, J. K., & Friedman, N. P. (2018). Rumination and Psychopathology: Are Anger and Depressive Rumination Differentially Associated with Internalizing and Externalizing Psychopathology? Clinical Psychological Science, 6(1), 18–31.

24. Emler, N. (1990). A Social Psychology of Reputation. European Review of Social Psychology, 1(1), 171–193.

25. Emler, N. (1994). Gossip, reputation, and social adaptation. In Good gossip (pp. 117–138). University Press of Kansas.

26. Epley, N., Keysar, B., Van Boven, L., & Gilovich, T. (2004). Perspective Taking as Egocentric Anchoring and Adjustment. Journal of Personality and Social Psychology, 87, 327–339.

27. Esteban, O., Blair, R., Markiewicz, C. J., Berleant, S. L., Moodie, C., Ma, F., Isik, A. I., Erramuzpe, A., Kent, M., Goncalves, M., & Others. (2018). fmriprep. Software: Practice & Experience.

28. Esteban, O., Markiewicz, C. J., Blair, R. W., Moodie, C. A., Isik, A. I., Erramuzpe, A., Kent, J. D., Goncalves, M., DuPre, E., Snyder, M., Oya, H., Ghosh, S. S., Wright, J., Durnez, J., Poldrack, R. A., & Gorgolewski, K. J. (2019). fMRIPrep: a robust preprocessing pipeline for functional MRI. Nature Methods, 16(1), 111–116.

29. Feurer, C., Jimmy, J., Chang, F., Langenecker, S. A., Phan, K. L., Ajilore, O., & Klumpp, H. (2021). Resting state functional connectivity correlates of rumination and worry in internalizing psychopathologies. Depression and Anxiety, 38(5), 488–497.

30. Finn, E. S., Corlett, P. R., Chen, G., Bandettini, P. A., & Constable, R. T. (2018). Trait paranoia shapes inter-subject synchrony in brain activity during an ambiguous social narrative. Nature Communications, 9(1), 2043.

31. Finn, E. S., Glerean, E., Khojandi, A. Y., Nielson, D., Molfese, P. J., Handwerker, D. A., & Bandettini, P. A. (2020). Idiosynchrony: From shared responses to individual differences during naturalistic neuroimaging. NeuroImage, 215, 116828.

32. Fossati, P., Hevenor, S. J., Graham, S. J., Grady, C., Keightley, M. L., Craik, F., & Mayberg, H. (2003). In search of the emotional self: an fMRI study using positive and negative emotional words. The American Journal of Psychiatry, 160(11), 1938–1945.

33. Fried, I., Mukamel, R., & Kreiman, G. (2011). Internally generated preactivation of single neurons in human medial frontal cortex predicts volition. Neuron, 69(3), 548–562.

34. Friston, K. (2018). Does predictive coding have a future? Nature Neuroscience, 21(8), 1019–1021.

35. Glasser, M. F., Sotiropoulos, S. N., Wilson, J. A., Coalson, T. S., Fischl, B., Andersson, J. L., Xu, J., Jbabdi, S., Webster, M., Polimeni, J. R., Van Essen, D. C., Jenkinson, M., & WU-Minn HCP Consortium. (2013). The minimal preprocessing pipelines for the Human Connectome Project. NeuroImage, 80, 105–124.

36. Goldberg, L. R. (1993a). The structure of personality traits: Vertical and horizontal aspects. In Studying lives through time: Personality and development (pp. 169–188). American Psychological Association.

37. Goldberg, L. R. (1993b). The Structure of Phenotypic Personality Traits. The American Psychologist, 9.

38. Gorgolewski, K., Burns, C. D., Madison, C., Clark, D., Halchenko, Y. O., Waskom, M. L., & Ghosh, S. S. (2011). Nipype: a flexible, lightweight and extensible neuroimaging data processing framework in python. Frontiers in Neuroinformatics, 5, 13.

39. Gorgolewski, K. J., Esteban, O., Markiewicz, C. J., Ziegler, E., Ellis, D. G., Notter, M. P., Jarecka, D., Johnson, H., Burns, C., Manhães-Savio, A., & Others. (2018). Nipype. Software: Practice & Experience.

40. Hastie, T., Tibshirani, R., & Friedman, J. (2008). The Elements of Statistical Learning: Data Mining, Inference, and Prediction (Second Edition). Springer.

41. Haxby, J. V., Connolly, A. C., & Guntupalli, J. S. (2014). Decoding neural representational spaces using multivariate pattern analysis. Annual Review of Neuroscience, 37(1), 435–456.

42. Hsieh, P.-J., Colas, J. T., & Kanwisher, N. G. (3/2012). Pre-stimulus pattern of activity in the fusiform face area predicts face percepts during binocular rivalry. Neuropsychologia, 50(4), 522–529.

43. Huang, Y.-F., Soon, C. S., Mullette-Gillman, O. A., & Hsieh, P.-J. (2014). Pre-existing brain states predict risky choices. NeuroImage, 101, 466–472.

44. Huang, Y., & Rao, R. P. N. (2011). Predictive coding. Wiley Interdisciplinary Reviews. Cognitive Science, 2(5), 580–593.

45. Hutchinson, J. B., & Barrett, L. F. (2019). The power of predictions: An emerging paradigm for psychological research. Current Directions in Psychological Science, 28(3), 280–291.

46. Inagaki, T. K., & Meyer, M. L. (2020). Individual differences in resting-state connectivity and giving social support: implications for health. Social Cognitive and Affective Neuroscience, 15(10), 1076–1085.

47. Ingram, R. E. (1990). Self-focused attention in clinical disorders: review and a conceptual model. Psychological Bulletin, 107(2), 156–176.

48. Ingram, R. E., & Smith, T. W. (1984). Depression and internal versus external focus of attention. Cognitive Therapy and Research, 8(2), 139–151.

49. Iyer, S., Collier, E., Broom, T. W., Finn, E. S., & Meyer, M. L. (2024). Individuals who see the good in the bad engage distinctive default network coordination during post-encoding rest. Proceedings of the National Academy of Sciences of the United States of America, 121(1), e2306295121.

50. Izuma, K., Kennedy, K., Fitzjohn, A., Sedikides, C., & Shibata, K. (2018). Neural Activity in the Reward-Related Brain Regions Predicts Implicit Self-Esteem: A Novel Validity Test of Psychological Measures Using Neuroimaging. Journal of Personality and Social Psychology. 10.1037/pspa0000114

51. Jacobson, N. S., & Anderson, E. A. (1982). Interpersonal skill and depression in college students: An analysis of the timing of self-disclosures. Behavior Therapy, 13(3), 271–282.

52. Javaheripour, N., Li, M., Chand, T., Krug, A., Kircher, T., Dannlowski, U., Nenadić, I., Hamilton, J. P., Sacchet, M. D., Gotlib, I. H., Walter, H., Frodl, T., Grimm, S., Harrison, B. J., Wolf, C. R., Olbrich, S., van Wingen, G., Pezawas, L., Parker, G., … Wagner, G. (2021). Altered resting-state functional connectome in major depressive disorder: a mega-analysis from the PsyMRI consortium. Translational Psychiatry, 11(1), 511.

53. Jenkins, A., & Mitchell, J. (2011). Medial prefrontal cortex subserves diverse forms of self-reflection. Social Neuroscience, 6, 211–218.

54. Just, N., & Alloy, L. B. (1997). The response styles theory of depression: tests and an extension of the theory. Journal of Abnormal Psychology, 106(2), 221–229.

55. Kaufman, G. F., & Libby, L. K. (2012). Changing beliefs and behavior through experience-taking. Journal of Personality and Social Psychology, 103(1), 1–19.

56. Kelley, W., Macrae, C., Wyland, C., Caglar, S., Inati, S., & Heatherton, T. (2002). Finding the self? An event-related fMRI Study. Journal of Cognitive Neuroscience, 14, 785–794.

57. Kragel, P. A., Koban, L., Barrett, L. F., & Wager, T. D. (2018). Representation, Pattern Information, and Brain Signatures: From Neurons to Neuroimaging. Neuron, 99(2), 257–273.

58. Kriegeskorte, N., Mur, M., & Bandettini, P. A. (2008). Representational similarity analysis - connecting the branches of systems neuroscience. Frontiers in Systems Neuroscience, 2. 10.3389/neuro.06.004.2008

59. Kucyi, A., Moayedi, M., Weissman-Fogel, I., Goldberg, M. B., Freeman, B. V., Tenenbaum, H. C., & Davis, K. D. (2014). Enhanced medial prefrontal-default mode network functional connectivity in chronic pain and its association with pain rumination. The Journal of Neuroscience: The Official Journal of the Society for Neuroscience, 34(11), 3969–3975.

60. Kuehner, C., & Weber, I. (1999). Responses to depression in unipolar depressed patients: an investigation of Nolen-Hoeksema’s response styles theory. Psychological Medicine, 29(6), 1323–1333.

61. Kutner, M. H., Nachtsheim, C., Neter, J., & Li, W. (2005). Applied Linear Statistical Models. McGraw-Hill Irwin.

62. Kwang, T., & Swann, W. B., Jr. (2010). Do people embrace praise even when they feel unworthy? A review of critical tests of self-enhancement versus self-verification. *Personality and Social Psychology Review: An Official Journal of the Society for Personality and Social Psychology*, Inc, 14(3), 263–280.

63. Landis, M. H., & Burtt, H. E. (1924). A Study of Conversations. Journal of Comparative Psychology, 4(1), 81–89.

64. Leary, M. R. (2007). Motivational and emotional aspects of the self. Annual Review of Psychology, 58(1), 317–344.

65. Libet, B., Gleason, C. A., Wright, E. W., & Pearl, D. K. (1983). Time of conscious intention to act in relation to onset of cerebral activity (readiness-potential). The unconscious initiation of a freely voluntary act. Brain: A Journal of Neurology, 106 (*Pt 3*), 623–642.

66. Lieberman, M. D., Jarcho, J. M., & Satpute, A. B. (2004). Evidence-based and intuition-based self-knowledge: an FMRI study. Journal of Personality and Social Psychology, 87(4), 421–435.

67. Lieberman, M. D., Straccia, M. A., Meyer, M. L., Du, M., & Tan, K. M. (04/2019). Social, self, (situational), and affective processes in medial prefrontal cortex (MPFC): Causal, multivariate, and reverse inference evidence. Neuroscience and Biobehavioral Reviews, 99, 311–328.

68. Liu, W., Kohn, N., & Fernández, G. (2019). Intersubject similarity of personality is associated with intersubject similarity of brain connectivity patterns. NeuroImage, 186, 56–69.

69. Mason, M. F., Norton, M. I., Van Horn, J. D., Wegner, D. M., Grafton, S. T., & Macrae, C. N. (2007). Wandering minds: the default network and stimulus-independent thought. Science, 315(5810), 393–395.

70. McKiernan, K. A., D’Angelo, B. R., Kaufman, J. N., & Binder, J. R. (2006). Interrupting the “stream of consciousness”: an fMRI investigation. NeuroImage, 29(4), 1185–1191.

71. Mehl, M. R. (2006). The lay assessment of subclinical depression in daily life. Psychological Assessment, 18(3), 340–345.

72. Meyer, M. L., Davachi, L., Ochsner, K. N., & Lieberman, M. D. (2019). Evidence That Default Network Connectivity During Rest Consolidates Social Information. Cerebral Cortex, 29(5), 1910–1920.

73. Meyer, M. L., & Lieberman, M. D. (2018). Why people are always thinking about themselves: Medial prefrontal cortex activity during rest primes self-referential processing. Journal of Cognitive Neuroscience, 30(5), 714–721.

74. Modi, S., Kumar, M., Kumar, P., & Khushu, S. (2015). Aberrant functional connectivity of resting state networks associated with trait anxiety. Psychiatry Research, 234(1), 25–34.

75. Murakami, M., Shteingart, H., Loewenstein, Y., & Mainen, Z. F. (2017). Distinct Sources of Deterministic and Stochastic Components of Action Timing Decisions in Rodent Frontal Cortex. Neuron, 94(4), 908–919.e7.

76. Naaman, M., Boase, J., & Lai, C.-H. (2010). Is it really about me? message content in social awareness streams. Proceedings of the 2010 ACM Conference on Computer Supported Cooperative Work, 189–192.

77. Nolen-Hoeksema, S. (1991). Responses to depression and their effects on the duration of depressive episodes. Journal of Abnormal Psychology, 100(4), 569–582.

78. Nolen-Hoeksema, S. (2000). The role of rumination in depressive disorders and mixed anxiety/depressive symptoms. Journal of Abnormal Psychology, 109(3), 504–511.

79. Nolen-Hoeksema, S., Parker, L. E., & Larson, J. (1994). Ruminative coping with depressed mood following loss. Journal of Personality and Social Psychology, 67(1), 92–104.

80. Park, H., & Rugg, M. D. (2010). Prestimulus hippocampal activity predicts later recollection. Hippocampus, 20(1), 24–28.

81. Poldrack, R. A. (2006). Can cognitive processes be inferred from neuroimaging data? Trends in Cognitive Sciences, 10(2), 59–63.

82. Qiu, Y., Wu, X., Liu, B., Huang, R., & Wu, H. (2024). Neural substrates of affective temperaments: An intersubject representational similarity analysis to resting-state functional magnetic resonance imaging in nonclinical subjects. Human Brain Mapping, 45(7), e26696.

83. Raichle, M. E. (2015). The Brain’s Default Mode Network. Annual Review of Neuroscience, 38(1), 433–447.

84. R Core Team. (2022). *R*: A Language and Environment for Statistical Computing. R Foundation for Statistical Computing. https://www.R-project.org/

85. Rogers, T. B., Kuiper, N., & Kirker, W. S. (1977). Self-reference and the encoding of personal information. Journal of Personality and Social Psychology. 10.1037//0022-3514.35.9.677

86. Ruby, F. J. M., Smallwood, J., Engen, H., & Singer, T. (2013). How self-generated thought shapes mood--the relation between mind-wandering and mood depends on the socio-temporal content of thoughts. PloS One, 8(10), e77554.

87. Sava-Segal, C., Richards, C., Leung, M., & Finn, E. S. (2023). Individual differences in neural event segmentation of continuous experiences. Cerebral Cortex, 33(13), 8164–8178.

87a. Scalabrini, A., Schimmenti, A., De Amicis, M., Porcelli, P., Benedetti, F., Mucci, C., & Northoff, G. (2022). The self and its internal thought: In search for a psychological baseline. Consciousness and Cognition, 97, 103244.

88. Schapiro, A. C., McDevitt, E. A., Rogers, T. T., Mednick, S. C., & Norman, K. A. (2018). Human hippocampal replay during rest prioritizes weakly learned information and predicts memory performance. Nature Communications, 9(1), 3920.

89. Schurger, A., Hu, P. ’ben’, Pak, J., & Roskies, A. L. (2021). What Is the Readiness Potential? Trends in Cognitive Sciences, 25(7), 558–570.

90. Schurger, A., Sitt, J. D., & Dehaene, S. (2012). An accumulator model for spontaneous neural activity prior to self-initiated movement. Proceedings of the National Academy of Sciences of the United States of America, 109(42), E2904–E2913.

91. Schwartz-Mette, R. A., & Rose, A. J. (2009). Conversational Self-Focus in Adolescent Friendships: Observational Assessment of an Interpersonal Process and Relations with Internalizing Symptoms and Friendship Quality. In Journal of Social and Clinical Psychology (Vol. 28, Issue 10, pp. 1263–1297). 10.1521/jscp.2009.28.10.1263

92. Sheline, Y. I., Barch, D. M., Price, J. L., Rundle, M. M., Vaishnavi, S. N., Snyder, A. Z., Mintun, M. A., Wang, S., Coalson, R. S., & Raichle, M. E. (2009). The default mode network and self-referential processes in depression. Proceedings of the National Academy of Sciences of the United States of America, 106(6), 1942–1947.

93. Smallwood, J., & Schooler, J. W. (2015). The Science of Mind Wandering: Empirically Navigating the Stream of Consciousness. Annual Review of Psychology, 66(1), 487–518.

94. Smith, T. W., & Greenberg, J. (1981). Depression and self-focused attention. Motivation and Emotion, 5(4), 323–331.

95. Song, X., & Wang, X. (2012). Mind wandering in Chinese daily lives--an experience sampling study. PloS One, 7(9), e44423.

96. Soon, C. S., Brass, M., Heinze, H.-J., & Haynes, J.-D. (2008). Unconscious determinants of free decisions in the human brain. Nature Neuroscience, 11(5), 543–545.

97. Soon, C. S., He, A. H., Bode, S., & Haynes, J.-D. (2013). Predicting free choices for abstract intentions. Proceedings of the National Academy of Sciences of the United States of America, 110(15), 6217–6222.

98. Spunt, B. (2016). *easy-optimize-x: Formal Release for Archiving on Zenodo*. Zenodo. https://zenodo.org/record/58616

99. Spunt, B., Meyer, M., & Lieberman, M. (2015). The Default Mode of Human Brain Function Primes the Intentional Stance. Journal of Cognitive Neuroscience, 27, 1–9.

100. Symons, C. S., & Johnson, B. T. (1997). The self-reference effect in memory: a meta-analysis. Psychological Bulletin, 121(3), 371–394.

101. Tambini, A., & Davachi, L. (2019). Awake Reactivation of Prior Experiences Consolidates Memories and Biases Cognition. Trends in Cognitive Sciences, 23(10), 876–890.

102. Tamir, D. I., & Mitchell, J. P. (2013). Anchoring and adjustment during social inferences. Journal of Experimental Psychology. General, 142(1), 151–162.

103. Tozzi, L., Zhang, X., Chesnut, M., Holt-Gosselin, B., Ramirez, C. A., & Williams, L. M. (2021). Reduced functional connectivity of default mode network subsystems in depression: Meta-analytic evidence and relationship with trait rumination. NeuroImage. Clinical, 30, 102570.

104. van Baar, J. M., Halpern, D. J., & FeldmanHall, O. (2021). Intolerance of uncertainty modulates brain-to-brain synchrony during politically polarized perception. Proceedings of the National Academy of Sciences of the United States of America, 118(20). 10.1073/pnas.2022491118

105. Van Essen, D. C., Smith, S. M., Barch, D. M., Behrens, T. E. J., Yacoub, E., Ugurbil, K., & WU-Minn HCP Consortium. (2013). The WU-Minn Human Connectome Project: an overview. NeuroImage, 80, 62–79.

106. Vu, H. L., Ng, K. T. W., Richter, A., & An, C. (2022). Analysis of input set characteristics and variances on k-fold cross validation for a Recurrent Neural Network model on waste disposal rate estimation. Journal of Environmental Management, 311, 114869.

107. Wager, T. D., Atlas, L. Y., Lindquist, M. A., Roy, M., Woo, C.-W., & Kross, E. (2013). An fMRI-based neurologic signature of physical pain. The New England Journal of Medicine, 368(15), 1388–1397.

108. Wagner, D. D., Chavez, R. S., & Broom, T. W. (2019). Decoding the neural representation of self and person knowledge with multivariate pattern analysis and data-driven approaches. Wiley Interdisciplinary Reviews. Cognitive Science, 10(1), e1482.

109. Wagner, D. D., Haxby, J. V., & Heatherton, T. F. (2012). The representation of self and person knowledge in the medial prefrontal cortex. Wiley Interdisciplinary Reviews. Cognitive Science, 3(4), 451–470.

110. Wallace, D. F. (2009). This Is Water: Some Thoughts, Delivered on a Significant Occasion, about Living a Compassionate Life (1st ed.). Little, Brown and Company.

111. Watkins, E. R. (2008). Constructive and unconstructive repetitive thought. Psychological Bulletin, 134(2), 163–206.

112. Watkins, E. R., & Roberts, H. (2020). Reflecting on rumination: Consequences, causes, mechanisms and treatment of rumination. Behaviour Research and Therapy, 127, 103573.

113. Woo, C.-W., Koban, L., Kross, E., Lindquist, M. A., Banich, M. T., Ruzic, L., Andrews-Hanna, J. R., & Wager, T. D. (2014). Separate neural representations for physical pain and social rejection. Nature Communications, 5(1), 5380.

114. Yeo, B. T. T., Krienen, F. M., Sepulcre, J., Sabuncu, M. R., Lashkari, D., Hollinshead, M., Roffman, J. L., Smoller, J. W., Zöllei, L., Polimeni, J. R., Fischl, B., Liu, H., & Buckner, R. L. (2011). The organization of the human cerebral cortex estimated by intrinsic functional connectivity. Journal of Neurophysiology, 106(3), 1125–1165.

115. Yu, H., Koban, L., Chang, L. J., Wagner, U., Krishnan, A., Vuilleumier, P., Zhou, X., & Wager, T. D. (2020). A Generalizable Multivariate Brain Pattern for Interpersonal Guilt. Cerebral Cortex, 30(6), 3558–3572.

116. Zhu, Y., Li, Z., Fan, J., & Han, S. (2007). Neural Basis of Cultural Influence on Self-Representation. NeuroImage, 34, 1310–1316.

