## Supplementary Materials for "A Neural Signature of the Bias Towards Self-Focus"

#### Study 1: Identifying a Neural Signature that Predicts Self-Focus

##### People consistently prefer to think about themselves over others, regardless of dimension and task timing

One possibility is that self-focus is driven by a particular dimension assessed, rather than a generalizable preference to think about oneself. For example, perhaps the desire to think about the self is driven specifically by participants' preference to think about their past. If so, then we would have identified a nuanced bias towards self-focus rather than a more general one. To assess this possibility, we tested whether target choice (who participants wanted to think about; self, friend, or Biden), as well as the response time to make the choice, interacted with the dimension considered (social roles, preferences, physical traits, personality traits, future, and past). First, we conducted a chi-square test of independence testing the interaction of the target choice and dimension. Then we ran an ANOVA test of the impact of dimension on response time. Target choice, but not response time, significantly interacted with dimensions (choice: $X^2$ (10, $N$ = 3447) = 112.01, $p$ < .001; response time: $F$(10, 3404.2) = 1.75, $p$ = .064, $\eta_p^2$ = 0.002). Because the target choice significantly interacted with the dimension, we followed up with separate paired sample t-tests for each target choice (self, friend, or Biden). These paired sample t-tests were performed using the R package *jmv.* Critically, these subsequent t-tests revealed that there was not a dimension within self choices that was significantly different than any of the others ($t$'s < 2.7; $p$'s > 0.50, bonferroni-corrected). Instead, the interaction was driven by the *other* target choices–friend choices and Biden choices. When participants selected to later think about their friend, they were more likely to choose to think about their friend's personality traits rather than social roles ($t$(31) = 4.20, $p$ = 0.009, Cohen's d = 0.87). When participants selected to think about Biden, they were more likely to choose to think about his social roles rather than preferences and past ($t_{preferences}$(31) = 4.00, $p$ = 0.017, Cohen's d = 0.87; $t_{past}$(31) = 3.50, $p$ = 0.045, Cohen's d = 0.82), and more likely to choose to think about his future rather than preferences ($t$(31) = 3.88, $p$ = 0.023, Cohen's d = 0.66). In other words, self-focus was not driven by a particular dimension(s), although when people preferred to think about their friend or Biden, certain dimensions were more preferred than others.

Another possibility is that the bias towards self-focus only occurs initially, in the beginning of the task, but this preference dissipates over time. This would suggest that people are not overwhelmingly self-focused, rather they can correct for this bias. To assess this possibility, binomial linear mixed models using the R package *lme4* were constructed to assess if there were changes in target choice over the course of the task (i.e., with time). Three models were constructed, one for each of the targets (self, friend, and Biden) with trial number, run number, and the interaction of trial number and run number included as regressors. There was no significant linear trend of choosing self over the length of the entire, two-run task ($\beta = 0.04$, standardized $\beta = 0.11$, $t(3447) = 0.40$, $p = 0.69$) nor for run one vs. run two ($\beta = -0.05$, standardized $\beta = 0.07$, $t(3447) = -0.67$, $p = 0.50$). Similarly, there was no significant choice by run interaction ($\beta = -0.01$, standardized $\beta = 0.07$, $t(3447) = -0.16$, $p = .87$). The same analyses for the other targets (i.e., friend; Biden) indicated that choice behavior also did not change with time for decisions to think about the friend (β's < .06; *p*'s > .60) or Biden (β's < .14; *p*'s > .14). Finally, we investigated whether the length of the jittered rest–which ranged from 2.5 to 6 seconds–impacted participants' choices of who they wanted to think about. If a longer jitter corresponded with a greater likelihood of choosing the self, it could indicate that active rehearsal during the brief rest (e.g. "next I want to think about myself") drove self-focused decisions as opposed to a more pre-reflective bias driven by spontaneous neural activity. Another binomial linear mixed effect model was constructed to assess if there were changes in target choice based on the length of the jittered rest that preceded the decision. However, the model showed no impact of jitter length on the subsequent trial's choice ($\beta = 0.006$, $t(3447) = 0.162$, $p = .87$). Collectively, the behavioral results suggest a persistent and spontaneous predisposition to quickly choose to focus on the self (vs. others).

**fMRI Preprocessing Pipeline**

For the fMRI dataset we collected, results included in this manuscript come from preprocessing performed using fMRIPrep 20.2.2 (Esteban et al., 2018, 2019) (RRID:SCR_016216), which is based on Nipype 1.6.1 (K. Gorgolewski et al., 2011; K. J. Gorgolewski et al., 2018) (RRID:SCR_002502). Per recommendations from the software developers, we report the exact text generated from the boilerplate below.

For each of the 2 BOLD runs found per subject (across all tasks and sessions), the following preprocessing was performed. First, a reference volume and its skull-stripped version were generated using a custom methodology of fMRIPrep. Head-motion parameters with respect to the BOLD reference (transformation matrices, and six corresponding rotation and translation parameters) are estimated before any spatiotemporal filtering using mcflirt (Jenkinson et al., 2002) (FSL 5.0.9). A B0-nonuniformity map (or fieldmap) was estimated based on a phase-difference map calculated with a dual-echo GRE (gradient-recall echo) sequence, processed with a custom workflow of SDCFlows inspired by the epidewarp.fsl script and further improvements in Human Connectome Project Pipelines (Glasser et al., 2013). The fieldmap was then co-registered to the target EPI (echo-planar imaging) reference run and converted to a displacements field map (amenable to registration tools such as ANTs) with FSL's fugue and other SDCflows tools. Based on the estimated susceptibility distortion, a corrected EPI (echo-planar imaging) reference was calculated for a more accurate co-registration with the anatomical reference. The BOLD time-series (including slice-timing correction when applied) were resampled onto their original, native space by applying a single, composite transform to correct for head-motion and susceptibility distortions. These resampled BOLD time-series will be referred to as preprocessed BOLD in original space, or just preprocessed BOLD. The BOLD reference was then co-registered to the T1w reference using flirt (Jenkinson & Smith, 2001) (FSL 5.0.9) with the boundary-based registration (Greve & Fischl, 2009) cost-function. Co-registration was configured with nine degrees of freedom to account for distortions remaining in the BOLD reference. Several confounding time-series were calculated based on the preprocessed BOLD: framewise displacement (FD), DVARS and three region-wise global signals. FD was computed using two formulations following Power (absolute sum of relative motions (2014)) and Jenkinson (relative root mean square displacement between affines (2002)). FD and DVARS are calculated for each functional run, both using their implementations in Nipype (following the definitions by Power et al. (2014)). The three global signals are extracted within the CSF, the WM, and the whole-brain masks. Additionally, a set of physiological regressors were extracted to allow for component-based noise correction (Behzadi et al., 2007) (CompCor). Principal components are estimated after high-pass filtering the preprocessed BOLD time-series (using a discrete cosine filter with 128s cut-off) for the two CompCor variants: temporal (tCompCor) and anatomical (aCompCor). tCompCor components are then calculated from the top 2% variable voxels within the brain mask. For aCompCor, three probabilistic masks (CSF, WM and combined CSF+WM) are generated in anatomical space. The implementation differs from that of Behzadi et al. (2007) in that instead of eroding the masks by 2 pixels on BOLD space, the aCompCor masks are subtracted from a mask of pixels that likely contain a volume fraction of GM. This mask is obtained by thresholding the corresponding partial volume map at 0.05, and it ensures components are not extracted from voxels containing a minimal fraction of GM. Finally, these masks are resampled into BOLD space and binarized by thresholding at 0.99 (as in the original implementation). Components are also calculated separately within the WM and CSF masks. For each CompCor decomposition, the k components with the largest singular values are retained, such that the retained components' time series are sufficient to explain 50 percent of variance across the nuisance mask (CSF, WM, combined, or temporal). The remaining components are dropped from consideration. The head-motion estimates calculated in the correction step were also placed within the corresponding confounds file. The confound time series derived from head motion estimates and global signals were expanded with the inclusion of temporal derivatives and quadratic terms for each (Satterthwaite et al., 2013). Frames that exceeded a threshold of 0.5 mm FD or 1.5 standardized DVARS were annotated as motion outliers. The BOLD time-series were resampled into standard space, generating a preprocessed BOLD run in MNI152NLin2009cAsym space. First, a reference volume and its skull-stripped version were generated using a custom methodology of fMRIPrep. All resamplings can be performed with a single interpolation step by composing all the pertinent transformations (i.e. head-motion transform matrices, susceptibility distortion correction when available, and co-registrations to anatomical and output spaces). Gridded (volumetric) resamplings were performed using antsApplyTransforms (ANTs), configured with Lanczos interpolation to minimize the smoothing effects of other kernels (Lanczos, 1964). Non-gridded (surface) resamplings were performed using mri_vol2surf (FreeSurfer).

**Parametric Modulation Methods**

Prior work examining pre-trial rest has used parametric modulation analyses to show that greater MPFC/BA10 activity during pre-trial rest accelerates self-reflection (Meyer & Lieberman, 2018). To see if we conceptually replicate this result in the main task, we performed a parametric modulation analysis, examining Yeo's MPFC ROI (2011) specifically, to determine if faster decisions to *choose* to think about the self are preferentially preceded by pre-trial MPFC/BA10 activity. We used *nltools* (Chang et al., 2023) in python to create a first-level model (performed on participants' residual images so that we can examine the jittered rest period without it being contaminated by task activation). The first-level model included three regressors reflecting the jittered rest period before each target choice: a rest before self-choice regressor, a rest before friend-choice regressor, and a rest before Biden-choice regressor. Each of these regressors had a parametric modulator regressor (orthogonalized with respect to that condition) that reflected the mean-centered response time for each trial in that regressor, yielding six total regressors for the experimental design. We created a contrast of self versus other–specifically comparing the parametric modulation regressors, where friend and Biden were combined into the other category. This contrast compared the speed to choose the self vs. the speed to choose others (friend and Biden). We focused on this self versus other contrast so that there were a near equal number of trials in each group (self = 1696, friend = 1127, Biden = 624, other = 1751) and to limit the number of comparisons we made. We then found the mean activation of this contrast in Yeo's MPFC ROI for each subject. Paired sample t-tests were performed on these results using the R package *stats* (R Core Team, 2022). A follow-up whole-brain parametric modulation analysis was completed using python's *nltools* (Chang et al., 2023) to investigate if any other brain regions demonstrated a parametric effect, and/or if other contrasts showed a parametric effect. We looked at four contrasts, 1) self vs other (friend and Biden), 2) self vs. friend, 3) self vs. Biden and 4) friend vs. Biden). We then performed a second-level model for each of these contrasts, utilizing *nltools* (Chang et al., 2023) in python, with a voxelwise false discovery rate corrected $p < .05$ threshold, contrast 1 k=99 cluster extent threshold, contrast 2 k=109 cluster extent threshold, contrast 3 k=49 cluster extent threshold, contrast 4 k=51 cluster extent threshold as determined by AFNI's 3dClustSim cluster size correction.

An additional follow-up whole-brain parcellation parametric modulation analysis was completed to investigate if any other brain regions defined by Yeo's (2011) 85 parcellation scheme demonstrated a parametric modulation effect, and/or if other contrasts showed a parametric modulation effect. We followed the same methodology of our primary MPFC/BA10 analysis. We used *nltools* (Chang et al., 2023) in python to create a first-level model (performed on participants' residual images) that included three regressors that captured the jittered rest period before each target choice: (1) a rest before self-choice regressor, (2) a rest before friend-choice regressor, and (3) a rest before Biden-choice regressor. Each of these regressors had a parametric modulator regressor (orthogonalized with respect to that condition) that reflected the mean-centered response time for each trial in that regressor, yielding six total regressors for the experimental design.. We looked at four contrasts of the parametric modulation regressors, 1) self vs other (friend and Biden), 2) self vs. friend, 3) self vs. Biden and 4) friend vs. Biden). We then found the mean activation of the contrasts in each of Yeo's 85 ROIs for each subject. Paired sample t-tests were performed on these results using the R package *stats* (R Core Team,

2022). We used a bonferroni corrected significance threshold of $p < .00059$ (.05/85) to account for multiple comparisons.

**The pre-self pattern temporally predicts the neural active self-reflection pattern during a resting state scan**

We investigated if the pre-self pattern could predict a neural marker of self-focus during a resting state scan in our sample of participants. To answer this question, we computed, for each subject, a multivariate pattern that reflected their active self-reflection in the self reflection fMRI task. We then used instatement analysis to test whether the presence of the pre-self pattern precedes the presence of this active self-reflection pattern in the core default network subsystem. This involved taking our pre-self pattern and each participant's multivariate active self-reflection pattern in the core subsystem and separately assessing each neural pattern's similarity with each second (i.e., TR) of the resting state scan. We then created linear mixed models with 0-20 TR delays: A) with the strength (i.e., correlation) of the pre-self pattern predicting the strength of the active self-reflection-pattern X TRs later and B) with the strength (i.e., correlation) of the active self-reflection pattern predicting the strength of the pre-self pattern X TRs later. The full results of this analysis are presented below in Supplemental Table 1, with all significant results (after Bonferroni correction) reported in the manuscripts results section.

**A. Pre-Self Predicting Active Self**

| TR delay | β | t-value | Uncorrected p-value |
|---|---|---|---|
| **0** | **-0.164** | **-4.70** | **<0.00001** |
| **1** | **-2.470** | **-7.16** | **<0.00001** |
| **2** | **-1.270** | **-3.65** | **0.00026** |
| 3 | 0.010 | 0.29 | 0.77300 |
| 4 | -0.070 | -1.98 | 0.04820 |
| 5 | 0.103 | 2.93 | 0.00341 |
| 6 | 0.101 | 2.87 | 0.00418 |
| 7 | 0.046 | 1.31 | 0.19100 |
| **8** | **0.162** | **4.55** | **<0.00001** |
| 9 | 0.108 | 3.04 | 0.00238 |
| 10 | 0.106 | 2.97 | 0.00299 |
| **11** | **0.160** | **4.44** | **0.00001** |
| 12 | 0.110 | 3.06 | 0.00226 |
| **13** | **0.143** | **3.94** | **0.00008** |
| **14** | **0.149** | **4.08** | **0.00005** |
| 15 | 0.106 | 2.89 | 0.00387 |
| 16 | 0.074 | 2.02 | 0.04370 |
| 17 | 0.085 | 2.30 | 0.02130 |
| 18 | 0.010 | 0.27 | 0.78600 |
| 19 | 0.037 | 0.98 | 0.32800 |
| 20 | 0.025 | 0.65 | 0.51400 |

**B. Active Self Predicting Pre-Self**

| TR delay | β | t-value | Uncorrected p-value |
|---|---|---|---|
| **0** | **-0.162** | **-4.70** | **<0.00001** |
| **1** | **-0.250** | **-7.24** | **<0.00001** |
| **2** | **-0.131** | **-3.79** | **0.00015** |
| 3 | 0.017 | 0.50 | 0.61900 |
| 4 | -0.083 | -2.37 | 0.01790 |
| 5 | 0.054 | 1.53 | 0.12500 |
| 6 | 0.037 | 1.05 | 0.29400 |
| 7 | -0.018 | -0.50 | 0.61800 |
| 8 | 0.072 | 2.03 | 0.04240 |
| 9 | 0.009 | 0.24 | 0.80700 |
| 10 | 0.005 | 0.14 | 0.88600 |
| 11 | 0.092 | 2.55 | 0.01070 |
| 12 | 0.002 | 0.05 | 0.95800 |
| 13 | 0.020 | 0.56 | 0.57300 |
| 14 | 0.057 | 1.58 | 0.11500 |
| 15 | -0.027 | -0.75 | 0.45600 |
| 16 | -0.043 | -1.18 | 0.23800 |
| 17 | -0.022 | -0.60 | 0.55000 |
| 18 | -0.033 | -0.89 | 0.37600 |
| 19 | 0.013 | 0.35 | 0.73000 |
| 20 | -0.040 | -1.09 | 0.27800 |

**Supplemental Table 1** To investigate the relationship between the presence of the pre-self pattern and the presence of the active self-reflection pattern, we ran linear mixed models with 0-20 TR delays: A) with the strength (i.e., correlation) of the pre-self pattern predicting the strength of the active self-reflection-pattern X TRs later and B) with the strength (i.e., correlation) of the active self-reflection pattern predicting the strength of the pre-self pattern X TRs later. To account for the number of statistical tests performed (41), we used a bonferroni-corrected significance value of $p$ < 0.00122 (i.e., .05x41). Tests that passed that bonferroni-corrected significance threshold are shown in bold.

**Study 2: Implications for Internalizing and Out-of-Sample Testing**

**The Anna Karenina Model results are not attributable to differences in pre-self pattern instatement variability**

One alternative explanation to the significant Anna Karenina results could be that people high on internalizing differ from others not on the temporal structure but instead on the strength or variability seen in their pre-self pattern instatement. We ran an additional analysis to investigate if the variability of subjects' pre-self pattern instatement was connected with their internalizing measure. To do this we took each subject's pre-self pattern instatement timecourse and calculated the standard deviation of those correlation values. Using a linear model, we found that there was no effect of internalizing on the standard deviation of participants' pre-self pattern instatement ($\beta$ = 0.00003, standardized $\beta$ = 0.01, $t(1085)$ = 0.44, and $p$ = 0.66).

**Testing an alternative internalizing "nearest neighbor" IS-RSA model in the Human Connectome Project Data**

To further confirm the Anna Karenina results are driven by the high internalizers, we ran a follow-up, complimentary IS-RSA referred to as a "Nearest Neighbor" model. The Nearest Neighbor model tests for similarity among pairs of participants who score similarly on a dimension (regardless of whether they score high or low on that dimension (Chang et al., 2020; Finn et al., 2020). Here, the Nearest Neighbor model tests whether low internalizers have a similar pre-self pattern timecourses to one another, as well as whether high internalizers have similar timecourses. In other words, each subject's timecourse is similar to other subjects' timecourses if they have a similar internalizing score, regardless of what the subjects' internalizing scores are.

Our Nearest Neighbor model represented the subject pair's dissimilarity of instatement vectors as the absolute value of the difference in the pair's internalizing rank. The less similar the pair's internalizing rank, the lower their similarity in pre-self pattern instatement timecourse, and the higher the similarity in the pair's internalizing rank, the higher their pre-self pattern instatement timecourse similarity. The Nearest Neighbor model was not significant ($r$ = -0.0003, $z$ = -0.14, $p$ = 0.89, Mantel permutation test). Additionally, none of the internalizing subscales showed a significant Nearest Neighbor effect (Anxiety and Depression: $r$ = -0.0008, $z$ = -0.34, $p$ = 0.73; Withdrawal: $r$ = 0.003, $z$ = 1.24, $p$ = 0.22; Somatic Complaints: $r$ = 0.003, $z$ = 1.28, $p$ = 0.20).

### References

Behzadi, Y., Restom, K., Liau, J., & Liu, T. T. (2007). A component based noise correction method (CompCor) for BOLD and perfusion based fMRI. *NeuroImage*, *37*(1), 90–101.

Chang, L., Jolly, E., Cheong, J. H., Burnashev, A., Chen, A., Clark, M., Frey, S., & Fitzpatrick, P. (2023). *cosanlab/nltools* (Version 0.5.0). https://doi.org/10.5281/zenodo.10059041

Chang, L., Manning, J., Baldassano, C., de la Vega, A., Fleetwood, G., Geerligs, L., Haxby, J., Lahnakoski, J., Parkinson, C., Shappell, H., Shim, W. M., Wager, T., Yarkoni, T., Yeshurun, Y., & Finn, E. (2020). *naturalistic-data-analysis/naturalistic_data_analysis: Version 1.0*. https://doi.org/10.5281/zenodo.3937849

Esteban, O., Blair, R., Markiewicz, C. J., Berleant, S. L., Moodie, C., Ma, F., Isik, A. I., Erramuzpe, A., Kent, M., Goncalves, M., & Others. (2018). fmriprep. *Software: Practice & Experience*.

Esteban, O., Markiewicz, C. J., Blair, R. W., Moodie, C. A., Isik, A. I., Erramuzpe, A., Kent, J. D., Goncalves, M., DuPre, E., Snyder, M., Oya, H., Ghosh, S. S., Wright, J., Durnez, J., Poldrack, R. A., & Gorgolewski, K. J. (2019). fMRIPrep: a robust preprocessing pipeline for functional MRI. *Nature Methods*, *16*(1), 111–116.

Finn, E. S., Glerean, E., Khojandi, A. Y., Nielson, D., Molfese, P. J., Handwerker, D. A., & Bandettini, P. A. (2020). Idiosynchrony: From shared responses to individual differences during naturalistic neuroimaging. *NeuroImage*, *215*, 116828.

Glasser, M. F., Sotiropoulos, S. N., Wilson, J. A., Coalson, T. S., Fischl, B., Andersson, J. L., Xu, J., Jbabdi, S., Webster, M., Polimeni, J. R., Van Essen, D. C., Jenkinson, M., & WU-Minn HCP Consortium. (2013). The minimal preprocessing pipelines for the Human Connectome Project. *NeuroImage*, *80*, 105–124.

Gorgolewski, K., Burns, C. D., Madison, C., Clark, D., Halchenko, Y. O., Waskom, M. L., & Ghosh, S. S. (2011). Nipype: a flexible, lightweight and extensible neuroimaging data processing framework in python. *Frontiers in Neuroinformatics*, *5*, 13.

Gorgolewski, K. J., Esteban, O., Markiewicz, C. J., Ziegler, E., Ellis, D. G., Notter, M. P., Jarecka, D., Johnson, H., Burns, C., Manhães-Savio, A., & Others. (2018). Nipype. *Software: Practice & Experience*.

Greve, D. N., & Fischl, B. (2009). Accurate and robust brain image alignment using boundary-based registration. *NeuroImage*, *48*(1), 63–72.

Jenkinson, M., Bannister, P., Brady, M., & Smith, S. (2002). Improved optimization for the robust and accurate linear registration and motion correction of brain images. *NeuroImage*, *17*(2), 825–841.

Jenkinson, M., & Smith, S. (2001). A global optimisation method for robust affine registration of brain images. *Medical Image Analysis*, *5*(2), 143–156.

Lanczos, C. (1964). Evaluation of Noisy Data. *Journal of the Society for Industrial and Applied Mathematics Series B Numerical Analysis*, *1*(1), 76–85.

Meyer, M. L., & Lieberman, M. D. (2018). Why people are always thinking about themselves: Medial prefrontal cortex activity during rest primes self-referential processing. *Journal of Cognitive Neuroscience*, *30*(5), 714–721.

Power, J. D., Mitra, A., Laumann, T. O., Snyder, A. Z., Schlaggar, B. L., & Petersen, S. E. (2014). Methods to detect, characterize, and remove motion artifact in resting state fMRI. *NeuroImage*, *84*, 320–341.

R Core Team. (2022). *R: A Language and Environment for Statistical Computing*. R Foundation for Statistical Computing. https://www.R-project.org/

Satterthwaite, T. D., Elliott, M. A., Gerraty, R. T., Ruparel, K., Loughead, J., Calkins, M. E., Eickhoff, S. B., Hakonarson, H., Gur, R. C., Gur, R. E., & Wolf, D. H. (2013). An improved framework for confound regression and filtering for control of motion artifact in the preprocessing of resting-state functional connectivity data. *NeuroImage*, *64*, 240–256.

Yeo, B. T. T., Krienen, F. M., Sepulcre, J., Sabuncu, M. R., Lashkari, D., Hollinshead, M.,

Roffman, J. L., Smoller, J. W., Zöllei, L., Polimeni, J. R., Fischl, B., Liu, H., & Buckner, R. L. (2011). The organization of the human cerebral cortex estimated by intrinsic functional connectivity. *Journal of Neurophysiology*, *106*(3), 1125–1165.
